# A switch in Nrg1-Rtg3 promoter recruitment coordinates mitochondrial cristae remodeling during the diauxic transition

**DOI:** 10.64898/2026.11.23.768758

**Authors:** Andrea de la Vega-Alvarado, Carlos Campero-Basaldua, Edgar Ramírez, Francisco Padilla-Garfias, Augusto César Poot-Hernández, Valeria Gómez-Marín, Ericka Moreno-Méndez, Ana Lilia Torres-Machorro, Sergio Encarnación-Guevara, Nayeli Torres-Ramírez, Rosario Ortíz-Hernández, Ximena Escalera-Fanjul, Héctor Quezada, James González, Lina Riego-Ruiz, Leonardo Peraza-Reyes, Alexander DeLuna, Claudio Scazzocchio, Alicia González

**Affiliations:** Departamento de Bioquímica y Biología Estructural, Instituto de Fisiología Celular, Universidad Nacional Autónoma de México, Mexico City, Mexico; Departamento de Biología Celular y Desarrollo, Instituto de Fisiología Celular, Universidad Nacional Autónoma de México, Mexico City, Mexico; Unidad de Bioinformática y Manejo de Información, Instituto de Fisiología Celular, Universidad Nacional Autónoma de México, Mexico City, Mexico; Centro de Investigación sobre el Envejecimiento, Centro de Investigación y de Estudios Avanzados del Instituto Politécnico Nacional (Cinvestav), Mexico City, Mexico; Laboratorio de Biología Celular, Departamento de Investigación en Fibrosis Pulmonar, Instituto Nacional de Enfermedades Respiratorias “Ismael Cosio Villegas”, Mexico City, Mexico; Centro de Ciencias Genómicas, Universidad Nacional Autónoma de México, Cuernavaca, Mexico; Departamento de Biología Celular, Facultad de Ciencias, Universidad Nacional Autónoma de México, Mexico City, Mexico; Laboratorio de Investigación en Inmunología y Proteómica, Hospital Infantil de México Federico Gómez, Mexico City, Mexico; División de Biología Molecular, Instituto Potosino de Investigación Científica y Tecnológica (IPICYT), San Luis Potosí, Mexico; Université Paris-Saclay, CEA, CNRS, Institute for Integrative Biology of the Cell (I2BC), 91198 Gif-sur-Yvette, France; Department of Life Sciences, Imperial College London, London, United Kingdom

**Keywords:** Transcriptional regulation, Transcriptional switch, Diauxic transition, Mitochondrial DNA, Mitochondrial nucleoid, Mitochondrial cristae, *Saccharomyces cerevisiae*

## Abstract

During the diauxic transition, *Saccharomyces cerevisiae* must preserve mitochondrial genome integrity while remodeling mitochondrial architecture for respiratory growth. The transcription factors Nrg1 and Rtg3 form a heteromeric regulator previously shown to be required for mitochondrial DNA maintenance, but its mitochondrial targets remained unknown. Here, we identify *ILV5* and *MGM101*, which encode mitochondrial nucleoid proteins, as Nrg1-Rtg3 targets. Both factors occupied these promoters, with Rtg3 binding independently and recruiting Nrg1, providing a direct regulatory link to mitochondrial DNA maintenance. Unexpectedly, electron microscopy revealed that loss of either factor caused severe cristae defects even during fermentative growth, and these defects were more pronounced after the diauxic transition. Nrg1-Rtg3 also switched from repressing *ATP20*, *ATP21*, and *MIC60* during fermentative growth to activating them after the transition. At the *ATP20* promoter, recruitment dependence also reversed across the diauxic transition, from Nrg1-dependent recruitment of Rtg3 during fermentation to Rtg3-dependent recruitment of Nrg1 after the transition. Even as this dependence reversed, both factors remained necessary to establish the wild-type chromatin organization associated with each transcriptional state. Together, these findings show how one transcriptional complex can maintain activation of mitochondrial nucleoid genes while switching the regulation of cristae-organizing genes as cells transition to respiratory growth.

## Introduction

The metabolic flexibility of *Saccharomyces cerevisiae* allows cells to favor fermentation when glucose is abundant while retaining the capacity to shift toward respiratory metabolism as fermentable carbon sources become limiting [1]. This diauxic transition requires extensive transcriptional reprogramming, including relief from glucose repression and activation of genes required for respiration, gluconeogenesis, and utilization of alternative carbon sources [2–4]. A central component of this response is the Snf1 kinase, which is activated as glucose declines and regulates several transcription factors, including the Mig1 repressor and the Cat8, Sip4, and Rds2 regulators of non-fermentative metabolism [3,5]. Adr1 and the Hap2/3/4/5 complex provide additional positive regulation of genes required for ethanol utilization and respiratory function [6,7]. Together, these pathways reorganize transcriptional control during the transition from fermentative to respiratory growth.

The diauxic transition also illustrates the regulatory challenge of how transcription factors change their output as cells shift between physiological states. Several factors involved in carbon-source adaptation act as both activators and repressors. Rds2 activates gluconeogenic genes while repressing genes that oppose gluconeogenesis [3,8], whereas Ert1 promotes genes associated with non-fermentative metabolism while repressing *PDC1*, which encodes a major pyruvate decarboxylase required for fermentation, during growth on ethanol [4]. These examples show that individual regulators can activate one set of genes while repressing another during carbon-source adaptation. A distinct regulatory problem arises when the same target switches between repression and activation as conditions change. Such reversals have been described in other signaling contexts, including the Sko1-Cyc8-Tup1 complex, which switches from repression to activation following Hog1 signaling in yeast [9], and *HapX-CBC*, which reverses its effect on the same target in response to iron availability in *Aspergillus* [10]. To our knowledge, however, a comparable promoter-bound switch by a heteromeric regulator has not been established across the *S. cerevisiae* diauxic transition. How heteromeric regulators reorganize promoter interactions to switch the same target genes between repression and activation therefore remains unclear.

Nrg1 and Rtg3 provide a relevant system to examine how a heteromeric regulator can support contrasting transcriptional outputs across metabolic states. Nrg1 contributes to glucose-dependent transcriptional repression through recruitment of the Cyc8-Tup1 corepressor complex, whereas Rtg3 functions with Rtg1 in the mitochondrial retrograde pathway and promotes expression of genes involved in metabolic adaptation and mitochondrial function [11,12]. Despite these distinct regulatory roles, both proteins are required for mtDNA maintenance, and loss of either causes severe mtDNA depletion and inability to sustain respiratory growth [13]. The transcriptional basis of this shared mitochondrial requirement remains unknown.

Respiratory adaptation also requires extensive remodeling of the mitochondrial inner membrane, where oxidative phosphorylation takes place. During the diauxic transition, mitochondria with relatively sparse cristae during fermentative growth develop more abundant and organized inner-membrane folds as respiratory capacity increases [14,15]. Cristae organization depends in part on ATP synthase oligomerization, promoted by ATP synthase subunits Atp20 and Atp21, and on the mitochondrial contact site and cristae organizing system, MICOS, whose central component Mic60 organizes crista junctions [15–17]. How transcriptional programs that preserve mitochondrial genome competence are integrated with remodeling of cristae architecture remains poorly understood.

Here, we show that Nrg1-Rtg3 directly activates *ILV5* and *MGM101*, genes encoding proteins associated with mitochondrial nucleoids, the mtDNA-containing structures that organize and maintain the mitochondrial genome [18]. This activation involves Rtg3-dependent recruitment of Nrg1 and changes in promoter nucleosome organization. Loss of either Nrg1 or Rtg3 also causes pronounced defects in mitochondrial cristae architecture. Nrg1-Rtg3 represses the cristae-organizing genes *ATP20*, *ATP21*, and *MIC60* during fermentation but activates them after the diauxic transition. Importantly, this switch coincides with a reversal in promoter recruitment, with Nrg1 recruiting Rtg3 during repression and Rtg3 recruiting Nrg1 during activation. In both regulatory states, however, both factors are required for WT chromatin organization. These findings identify reciprocal promoter recruitment between mutually dependent Nrg1 and Rtg3 as a mechanism linking mitochondrial genome maintenance with transcriptional control of cristae remodeling.

## Results

### Nrg1-Rtg3 directly activates mitochondrial nucleoid genes through Rtg3-dependent promoter recruitment

Previous work showed that the Nrg1-Rtg3 regulator is required for mitochondrial DNA maintenance, as loss of either factor results in mtDNA depletion and loss of respiratory competence. To identify transcriptional targets that could account for this phenotype, we carried out RNA-seq comparing untreated wild-type (WT) cells with alanine-treated WT, *nrg1*Δ, and *rtg3*Δ cells, under conditions previously associated with Nrg1-Rtg3 activity [13] (**Fig. 1A**; **Dataset S1**). Principal-component analysis separated untreated WT cells from the alanine-treated samples and further distinguished alanine-treated WT cells from both mutants, indicating that loss of either factor substantially alters the transcriptional response to alanine. The analysis revealed a broad bidirectional response, with 1,057 genes showing an activation pattern dependent on both factors and 926 showing the reciprocal repression pattern. Consistent with this classification, comparison of gene responses in WT and the two mutants revealed large groups of genes whose alanine-dependent expression changes were attenuated or enhanced in the absence of Nrg1 or Rtg3 (**Fig. 1B**). The former were enriched for functions including translation, ribosome biogenesis, DNA replication, and cell-cycle progression, while the latter were enriched for autophagy, proteolysis, lipid and energy metabolism, and peroxisomal processes (**Fig. S1**).

**Figure 1.**
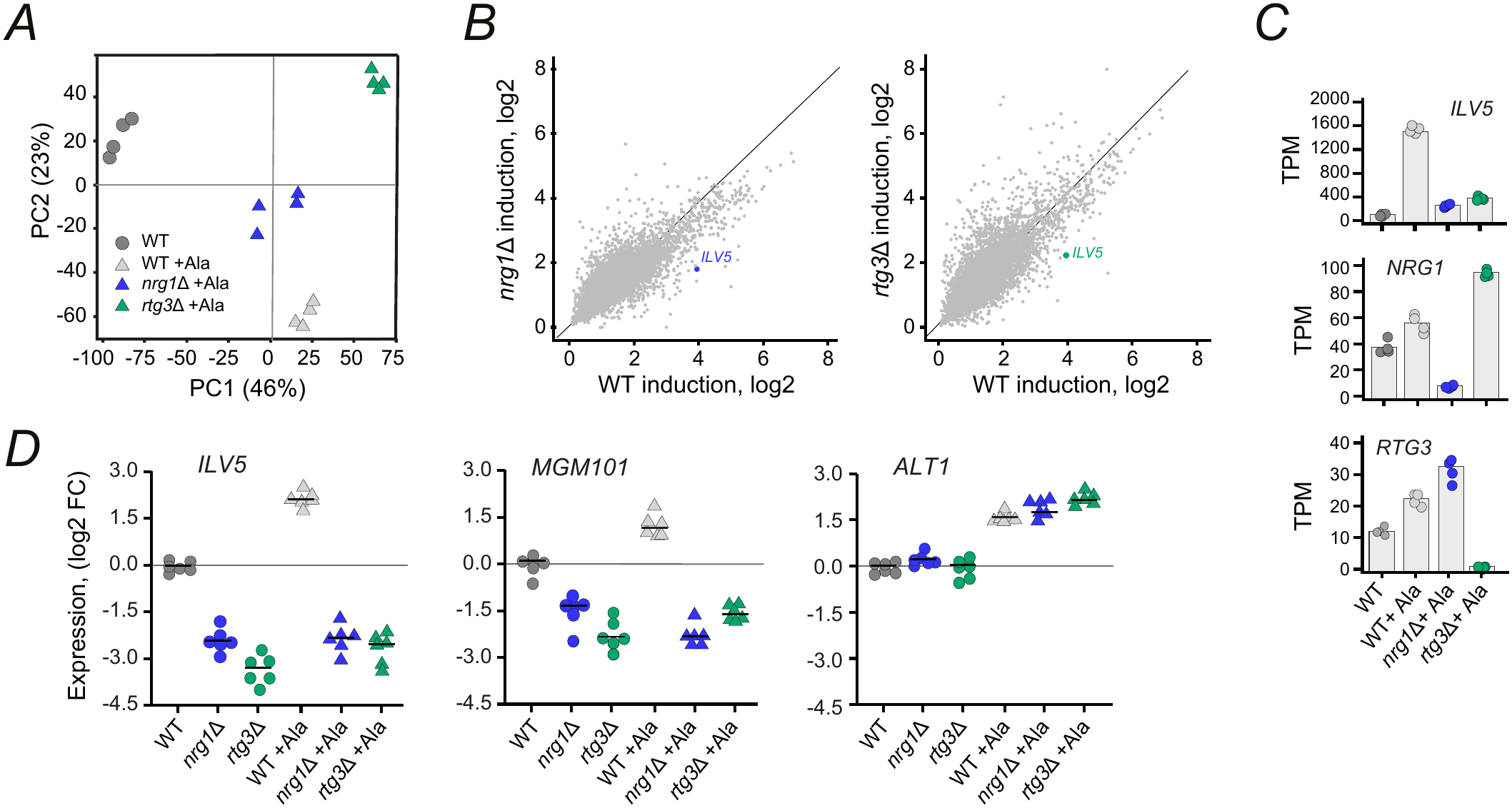
Alanine-responsive expression identifies Nrg1-Rtg3-dependent targets. **(A)** Principal component analysis (PCA) of RNA-seq profiles from WT cells grown in minimal medium containing GABA (7 mM), and WT, *nrg1*Δ, and *rtg3*Δ cells grown in GABA plus alanine (7 mM). **(B)** Comparison of alanine-induced log2 expression changes in WT cells (horizontal axis) with those in *nrg1*Δ or *rtg3*Δ cells (vertical axis). *ILV5* is indicated. **(C)** RNA-seq expression levels (transcripts per million) of *ILV5*, *NRG1*, and *RTG3* in the indicated strains and conditions. **(D)** RT-qPCR analysis of *ILV5*, *MGM101*, and *ALT1* in WT, *nrg1*Δ, and *rtg3*Δ cells grown with γ-aminobutyric acid (GABA) as sole nitrogen source or GABA plus alanine. Expression was normalized to 18S rRNA and calculated by the 2^−ΔΔCt^ method. Individual symbols represent biological replicates; horizontal lines indicate the mean.

Within this extensive regulatory landscape, the previously described mtDNA phenotype pointed to a particularly relevant subset of Nrg1-Rtg3 targets. *ILV5*, a metabolic enzyme with an established moonlighting role in mitochondrial nucleoid organization and mtDNA stability [19,20], showed Nrg1-and Rtg3-dependent activation **(Fig. 1B–D)**. Its strong induction by alanine in WT cells was markedly decreased in both *nrg1*Δ and *rtg3*Δ cells in the RNA-seq data and independently confirmed by RT-qPCR **(Fig. 1C,D)**. We therefore examined additional nucleoid-associated genes and identified *MGM101*, a central factor in mitochondrial genome maintenance, with a similar regulatory signature (**Fig. 1D**; **Fig. S2**). Other nucleoid genes did not show this concordant regulatory pattern; *ACO1* changed in opposite directions in the two mutants, *ABF2* increased predominantly in *nrg1*Δ cells, and *RIM1* was little affected by alanine in WT cells and selectively reduced in *rtg3*Δ, indicating dependence on Rtg3 but not Nrg1 (**Fig. S2**). Thus, among the nucleoid-associated genes examined, *ILV5* and *MGM101* showed the clearest concordant signature of alanine-responsive activation requiring both Nrg1 and Rtg3 and were selected for further analysis. As a control for general alanine responsiveness, *ALT1* induction was maintained in both mutants, excluding a general defect in transcriptional response to alanine (**Fig. 1D**). RNA-seq experiments also confirmed loss of the cognate *NRG1* or *RTG3* transcript in each deletion strain, while expression of the other factor was retained, excluding reduced expression of one factor as a secondary consequence of deleting the other (**Fig. 1C**).

The similar transcriptional functions of Nrg1 and Rtg3 were consistent with a coordinated action in terms of their physical interaction. Co-immunoprecipitation of epitope-tagged Nrg1 and Rtg3 detected their association with either tag-specific antibody, whereas no corresponding signal was recovered with the HA (hemagglutinin) control; the interaction was reproduced in three independent biological replicates (**Fig. 2A**). Thus, Nrg1 and Rtg3 physically associate under the same alanine-responsive conditions in which they coordinately promote *ILV5* and *MGM101* expression. Together, these observations focused our analysis on *ILV5* and *MGM101* as candidate targets linking Nrg1-Rtg3 activity to mitochondrial genome maintenance.

**Figure 2.**
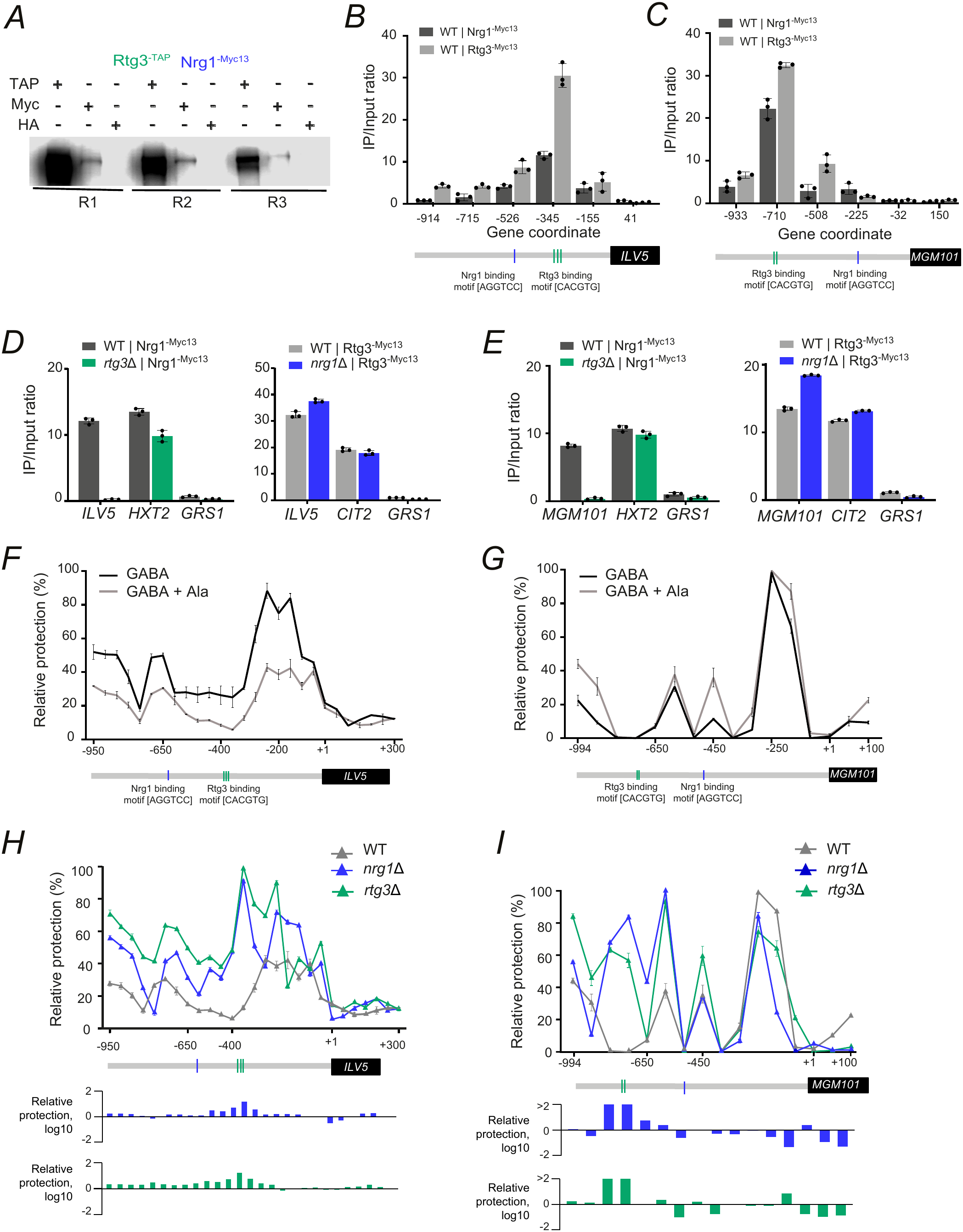
Nrg1-Rtg3 promoter recruitment and chromatin organization at the mitochondrial nucleoid genes *ILV5* and *MGM101*. **(A)** Co-immunoprecipitation analysis of Nrg1-Myc13 and Rtg3-TAP in three independent replicates (R1–R3); HA was included as a negative control. **(B,C)** ChIP-qPCR profiles of Nrg1-Myc13 and Rtg3-Myc13 across the *ILV5* **(B)** and *MGM101* **(C)** promoter regions. Amplicon coordinates are relative to the translation start site; predicted Rtg3-binding sites are indicated in green. **(D,E)** ChIP-qPCR of Nrg1-Myc13 in WT and *rtg3*Δ cells and Rtg3-Myc13 in WT and *nrg1*Δ cells grown on GABA plus alanine at *ILV5* **(D)** and *MGM101* **(E)**. *HXT2* and *CIT2* were included as positive controls for Nrg1 and Rtg3 binding, respectively, with *GRS1* as a negative control. ChIP enrichment is shown as the IP/input ratio. **(F,G)** Nucleosome scanning analysis of the *ILV5* **(F)** and *MGM101* **(G)** promoters in WT cells grown in minimal medium containing glucose (2%) and GABA (7 mM), with or without alanine (7 mM). Cells were collected at OD_600_ = 0.6 and analyzed as described in Methods. Relative protection is shown across each promoter. Predicted Rtg3-binding sites are indicated in green and predicted Nrg1-binding sites in blue. Data are shown as mean ± SEM from three independent experiments. **(H,I)** Nucleosome scanning analysis across the *ILV5* **(H)** and *MGM101* **(I)** promoters in WT, *nrg1*Δ, and *rtg3*Δ cells grown in minimal medium containing glucose (2%), GABA (7 mM), and alanine (7 mM). Upper plots show relative protection across each promoter; lower plots show the corresponding log10 relative protection values for *nrg1*Δ and *rtg3*Δ. Predicted Rtg3-binding sites are indicated in green. ChIP-qPCR and nucleosome-scanning data represent three independent experiments.

We next asked whether Nrg1-Rtg3 directly regulated *ILV5* and *MGM101*. ChIP analysis showed that both Rtg3 and Nrg1 were enriched locally at the two promoters (**Fig. 2B,C**). At *ILV5*, the strongest Rtg3 enrichment occurred around the region containing the cluster of annotated Rtg3-binding motifs, and Nrg1 showed a coincident peak despite the annotated Nrg1-binding motif lying farther upstream (**Fig. 2B**). A similar arrangement was observed at *MGM101*, with both factors strongly enriched around the upstream Rtg3-binding site, whereas occupancy was substantially lower around the downstream Nrg1-binding motif (**Fig. 2C**). Thus, at both promoters, Nrg1 occupancy overlapped with Rtg3 binding rather than with the position of the predicted Nrg1-binding site. This recruitment architecture was different from that previously observed at the Nrg1-Rtg3-repressed *ALT2* promoter, where Nrg1 can bind independently of Rtg3 [13]. At both *ILV5* and *MGM101*, Nrg1 occupancy was abolished in *rtg3*Δ cells, whereas Rtg3 binding was retained in *nrg1*Δ cells (**Fig. 2D,E**). Moreover, Nrg1 occupancy at the control *HXT2* promoter remained detectable in *rtg3*Δ cells, while Rtg3 occupancy at *CIT2* was maintained in *nrg1*Δ cells; binding to the negative-control *GRS1* locus remained near background in all cases (**Fig. 2D,E**). These controls indicate that the loss of Nrg1 recruitment at *ILV5* and *MGM101* is promoter-specific and does not reflect a general defect in Nrg1 abundance or ChIP recovery.

The close correspondence between Nrg1-Rtg3 recruitment and transcriptional activation prompted us to examine whether both factors also shape local nucleosome organization at these promoters. In WT cells, alanine markedly reduced relative protection across much of the *ILV5* promoter (**Fig. 2F**), consistent with increased chromatin accessibility, whereas the *MGM101* promoter showed little overall change (**Fig. 2G**). Further nucleosome scanning under inducing conditions revealed substantially lower relative protection across much of the upstream *ILV5* promoter in WT cells than in either mutant, with the strongest difference centered on the region containing the Rtg3-binding sites (**Fig. 2H**). Deletion of either *NRG1* or *RTG3* markedly increased protection across this interval, consistent with increased nucleosome occupancy over a region that is comparatively accessible in WT cells. The *MGM101* promoter also showed pronounced Nrg1-and Rtg3-dependent changes, consistent with nucleosome redistribution (**Fig. 2I**). In particular, both mutants gained protection across the upstream region encompassing the Rtg3-binding site while showing reduced protection at several more promoter-proximal positions. The parallel changes produced by the two deletions at both promoters further indicate that Nrg1 and Rtg3 act together to maintain the WT chromatin configuration. Thus, although Rtg3 binds independently and recruits Nrg1, both factors are required to maintain WT promoter nucleosome organization at *ILV5* and *MGM101* under inducing conditions.

To determine whether the same recruitment relationship extended to a nucleoid gene that did not require both factors for expression, we analyzed Nrg1 and Rtg3 occupancy at the *RIM1* promoter. In contrast to *ILV5* and *MGM101*, both factors remained promoter-bound in the absence of the other, indicating largely independent recruitment at *RIM1* (**Fig. S3**). This difference was also reflected functionally, as *RIM1* expression was strongly reduced in *rtg3*Δ but largely maintained in *nrg1*Δ cells, and nucleosome organization was altered much more extensively in *rtg3*Δ than in *nrg1*Δ cells. Thus, *RIM1* is regulated predominantly by Rtg3 and does not display the Rtg3-dependent recruitment of Nrg1 observed at *ILV5* and *MGM101*. These results indicate that Rtg3-dependent Nrg1 recruitment is not a general feature of nucleoid-gene promoters but is associated with the concordant Nrg1-Rtg3 regulation of *ILV5* and *MGM101*.

If *ILV5* and *MGM101* are functionally relevant outputs of Nrg1-Rtg3 regulation, their loss should recapitulate at least some of the mitochondrial alterations caused by loss of Nrg1 or Rtg3. Because Ilv5 and Mgm101 have established roles in mitochondrial genome maintenance [19,21,22], we asked whether the mitochondrial defects caused by their loss recapitulated those caused by loss of Nrg1 or Rtg3. Specifically, we compared respiratory competence and mitochondrial genome maintenance in *nrg1*Δ, *rtg3*Δ, *ilv5*Δ, and *mgm101*Δ cells. All four mutants remained capable of growth under fermentative conditions on glucose, although *rtg3*Δ and *mgm101*Δ showed reduced growth rates relative to WT, with a more modest reduction in *nrg1*Δ (**Fig. 3A**). The distinction between strains became much more pronounced when ethanol was provided as the sole carbon source, under conditions in which respiratory metabolism was required. We observed that *nrg1Δ* and *rtg3Δ* cells showed no detectable growth on ethanol (**Fig. 3B**). The *mgm101Δ* mutant showed a similarly severe respiratory defect, whereas *ilv5Δ* cells retained growth on ethanol, although at approximately half the WT growth rate. These differences closely corresponded to mitochondrial genome content; measurement of three independent mtDNA loci, *COX2*, *COX3*, and *ATP6*, showed mtDNA retention across all three loci in WT cells but no detectable mtDNA in *nrg1*Δ, *rtg3*Δ, or *mgm101*Δ cells. In contrast, *ilv5*Δ retained approximately half of the WT mtDNA level at each locus (**Fig. 3C**). Respiratory growth closely corresponded with mtDNA retention across strains, whereby mutants with essentially complete mtDNA loss were unable to grow on ethanol, whereas partial mtDNA retention in *ilv5*Δ was accompanied by residual respiratory growth.

**Figure 3.**
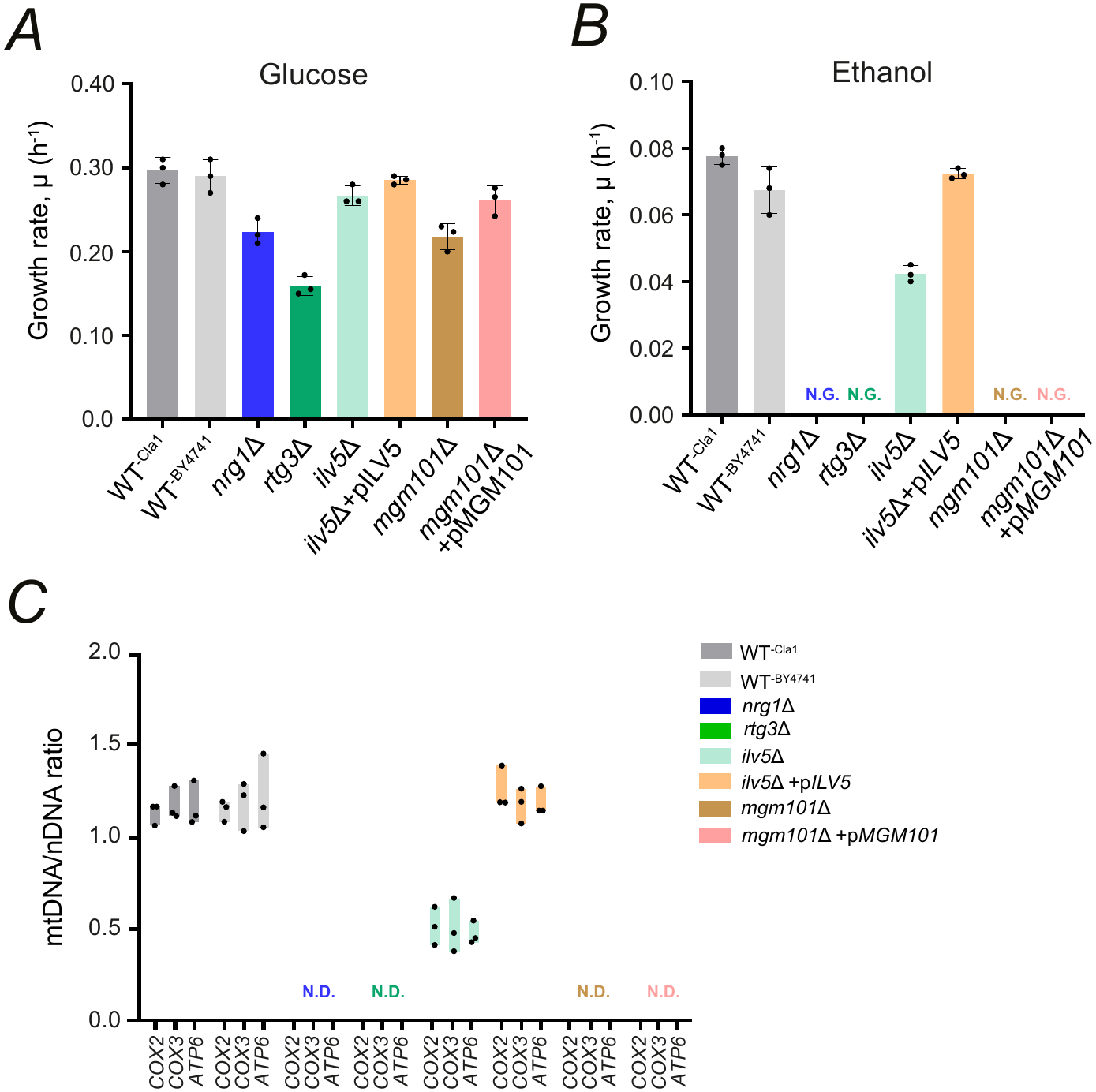
Loss of Nrg1 or Rtg3 compromises respiratory growth and mtDNA maintenance. (A,B) Specific growth rates of the indicated strains in minimal medium containing 40 mM ammonium sulfate and either 2% glucose **(A)** or 2% ethanol **(B)**. Data are from three independent experiments. N.G., no growth. **(C)** Relative mtDNA abundance determined by qPCR using the mitochondrial loci *COX2*, *COX3*, and *ATP6* relative to the nuclear *COX6* locus. N.D., not detectable. CLA1 and BY4741 were the WT parental backgrounds for the corresponding mutants; *ilv5Δ* and *mgm101Δ* strains carrying centromeric plasmids with the cognate WT genes are indicated as +p*ILV5* and +p*MGM101*, respectively.

RT-qPCR analysis of additional mitochondrial genes showed no detectable expression of *COX2*, *COX3*, *ATP6*, *ATP9*, or *COB1* in *nrg1*Δ, *rtg3*Δ, or *mgm101*Δ cells, whereas expression was retained in *ilv5*Δ cells. Expression of the nuclear-encoded mitochondrial gene *COX6* was maintained in all strains, indicating a selective defect in mitochondrial gene expression (**Fig. S4**). To determine whether the association between mtDNA loss and respiratory deficiency extended to other mitochondrial nucleoid components, we examined *abf2*Δ, *aco1*Δ, and *rim1*Δ cells. All three mutants showed no detectable mtDNA across the *COX2*, *COX3*, and *ATP6* loci and failed to grow on ethanol, despite retaining growth on glucose, although at different rates (**Fig. S4**). Thus, disruption of multiple mitochondrial nucleoid components similarly couples mtDNA loss to loss of respiratory competence.

Together, these results establish *ILV5* and *MGM101* as direct Nrg1-Rtg3 targets, with Rtg3-dependent recruitment of Nrg1 associated with promoter chromatin organization and transcriptional activation. Because both genes encode mitochondrial nucleoid proteins and their disruption compromises mtDNA maintenance and respiratory growth, their regulation links Nrg1-Rtg3 activity to mitochondrial nucleoid function.

### Loss of Nrg1-Rtg3 function couples mitochondrial genome defects to cristae-deficient architecture across the diauxic transition

The similarity between the Nrg1-Rtg3 phenotype and that produced by disruption of multiple mitochondrial nucleoid components prompted us to ask whether loss of Nrg1 or Rtg3 affected mitochondrial organization beyond mtDNA maintenance. This question was particularly relevant across the diauxic transition, when *S. cerevisiae* shifts from predominantly fermentative to respiratory metabolism and mitochondria normally undergo extensive structural remodeling, including a marked increase in cristae abundance and organization [15].

We therefore examined mitochondrial ultrastructure by transmission electron microscopy in wild-type, *nrg1*Δ, and *rtg3*Δ cells during exponential growth and after the diauxic transition (**Fig. 4**). Representative sections revealed a pronounced disruption of mitochondrial internal architecture in both mutants. During exponential growth, WT cell sections were dominated by mitochondria with recognizable, regularly organized cristae, whereas normal mitochondrial profiles were scarce in both *nrg1*Δ and *rtg3*Δ cells (**Fig. 4A**). Quantification of all cell sections confirmed a shift toward abnormal morphologies, with *nrg1*Δ particularly enriched for mitochondria lacking recognizable cristae, together with irregular mitochondria, whereas *rtg3*Δ displayed a broader distribution of irregular, cristae-deficient, and mixed defects, with occasional onion-ring structures (**Fig. 4B**). The complete phenotype distributions and underlying cell counts for WT, *nrg1*Δ, and *rtg3*Δ in both growth phases are provided in **Fig. S5**. Thus, loss of either factor strongly shifted the population away from normal mitochondrial architecture.

**Figure 4.**
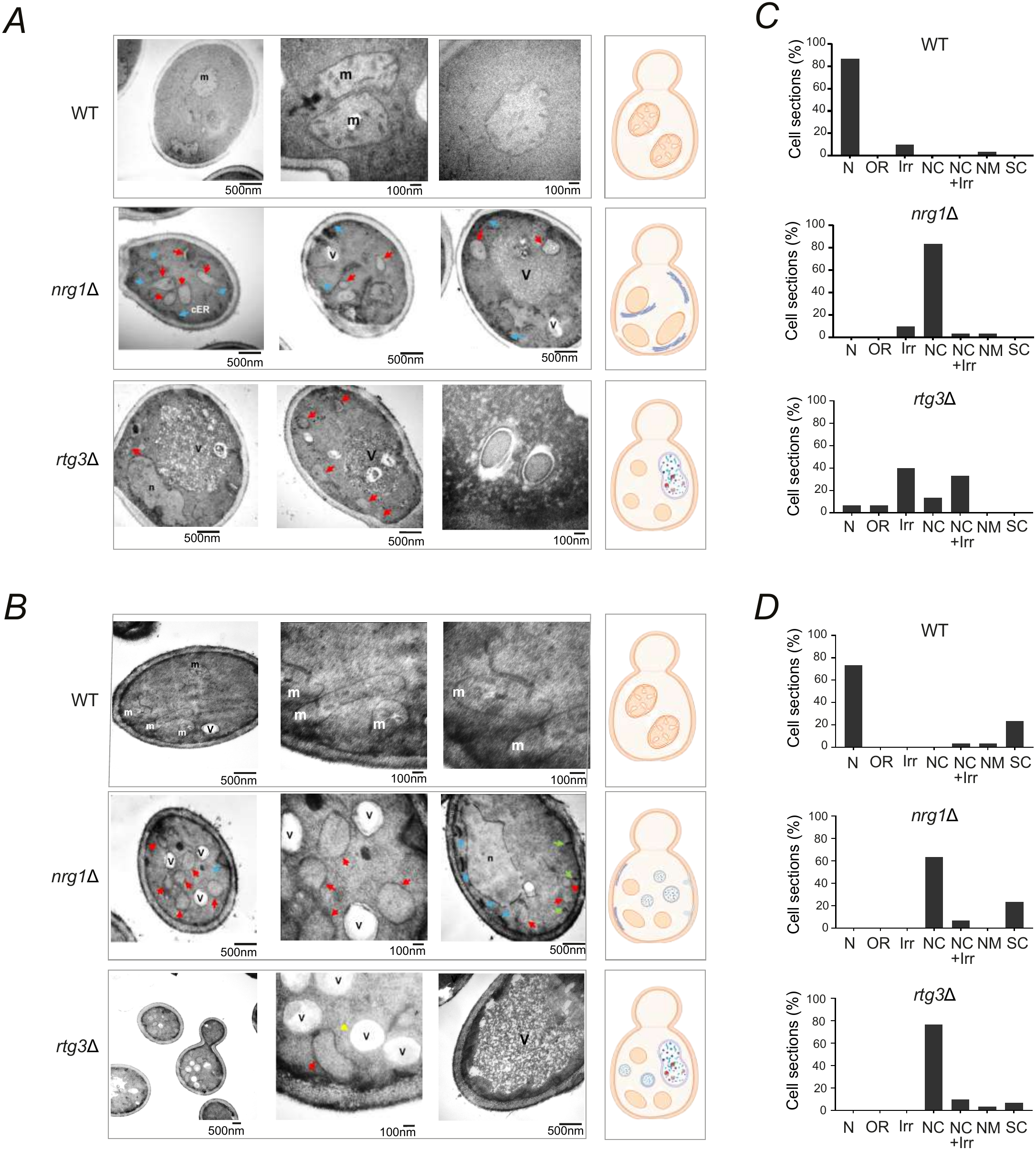
Loss of Nrg1 or Rtg3 reveals cristae-deficient mitochondrial architecture across the diauxic transition. (A,C) Three representative TEM images are shown for each strain, together with a schematic on the right summarizing the most common mitochondrial morphology observed, during exponential growth in 2% glucose **(A)** and after ∼25 h of growth in the same medium, following the diauxic transition (OD_600_ = 1.0) **(C)**. Red arrows indicate cristae-deficient mitochondrial profiles; blue arrows indicate putative cortical ER, green arrows indicate plasma-membrane invaginations, and the yellow arrow indicates a putative double-membrane autophagosome. **n**, nucleus; **m**, mitochondrion; **V**, vacuole; **cER**, cortical endoplasmic reticulum. Scale bars are indicated. **(B,D)** Quantification of mitochondrial phenotypes in 30–40 cell sections examined during exponential **(B)** and post-diauxic **(D)** growth. N, normal; OR, onion-ring structures; Irr, irregular mitochondria; NC, mitochondria lacking cristae; NC+Irr, combined no-cristae and irregular phenotypes; NM, no mitochondria observed; SC, short cristae.

The contrast between WT and the two mutants became particularly evident after the diauxic transition, when WT mitochondria had developed the more elaborate cristae architecture associated with respiratory growth, whereas both *nrg1*Δ and *rtg3*Δ cells continued to display predominantly abnormal mitochondrial profiles. Normal mitochondrial profiles predominated in WT cells but remained scarce in both mutants, where irregular, cristae-deficient, and mixed morphologies continued to predominate (**Fig. 4C,D**). The persistence of mitochondria lacking recognizable cristae was particularly evident in both mutants under a metabolic condition in which WT mitochondria showed increased cristae development. The persistence of these phenotypes across both growth phases indicates that defective cristae architecture is already established during fermentative growth and is not restored during the subsequent transition to respiration. Together, these results reveal that the mitochondrial consequences of loss of Nrg1 or Rtg3 extend beyond genome maintenance and respiratory competence to a persistent defect in cristae remodeling across the diauxic transition.

### Nrg1-Rtg3 switches from repression to activation of cristae-organizing genes during the diauxic transition

The pronounced cristae defects in *nrg1*Δ and *rtg3*Δ cells suggested that Nrg1-Rtg3 regulates not only mitochondrial genome maintenance but also genes more directly involved in inner-membrane organization. We therefore examined *ATP20*, *ATP21*, and *MIC60*, which encode key determinants of cristae architecture. *ATP20* and *ATP21* encode ATP synthase subunits required for dimerization and normal cristae morphology, whereas *MIC60* encodes a core component of the MICOS complex that organizes crista junctions [16,17]. To determine whether their regulation changes across metabolic states, we measured expression of these genes by RT-qPCR in WT, *nrg1*Δ, and *rtg3*Δ cells during fermentative and post-diauxic growth in GABA medium with or without alanine.

The three cristae-organizing genes showed a coordinated switch in the effect of Nrg1 or Rtg3 loss across the diauxic transition. During fermentative growth, *ATP20*, *ATP21*, and *MIC60* were expressed at higher levels in *nrg1*Δ and *rtg3*Δ cells than in WT, indicating repression by Nrg1-Rtg3 under these conditions (**Fig. 5A–C**). After the diauxic transition, expression decreased in both mutants and fell well below WT levels, indicating that Nrg1 and Rtg3 are required to maintain expression of all three genes in the post-diauxic state. Thus, the defining feature is a reversal in the direction of the mutant effect relative to WT, from increased expression during fermentation to strongly reduced expression after the diauxic transition. *ILV5* provided an important contrast. Although its expression also decreased in the mutants after the diauxic transition, it was already below WT during fermentative growth and remained below WT in the post-diauxic state, indicating continued positive regulation by Nrg1 and Rtg3 across both metabolic states and no switch from repression to activation (**Fig. 5D**). The control gene *ALT1* likewise showed no comparable reversal (**Fig. 5E**). These patterns were observed both with GABA as the sole nitrogen source and with GABA supplemented with alanine, indicating that alanine modulates expression within each state without determining the switch from repression to activation. The contrasting behavior of the cristae-organizing genes, *ILV5*, and *ALT1* indicates that the diauxic transition reshapes Nrg1-Rtg3 regulatory output in a target-specific manner. Importantly, the genes undergoing the repression-to-activation switch encode key determinants of inner-membrane architecture. Their altered regulation therefore provides a transcriptional link between Nrg1-Rtg3 and the cristae defects observed in *nrg1*Δ and *rtg3*Δ cells (**Fig. 4**), and a plausible basis for their failure to establish normal cristae organization across the diauxic transition.

**Figure 5.**
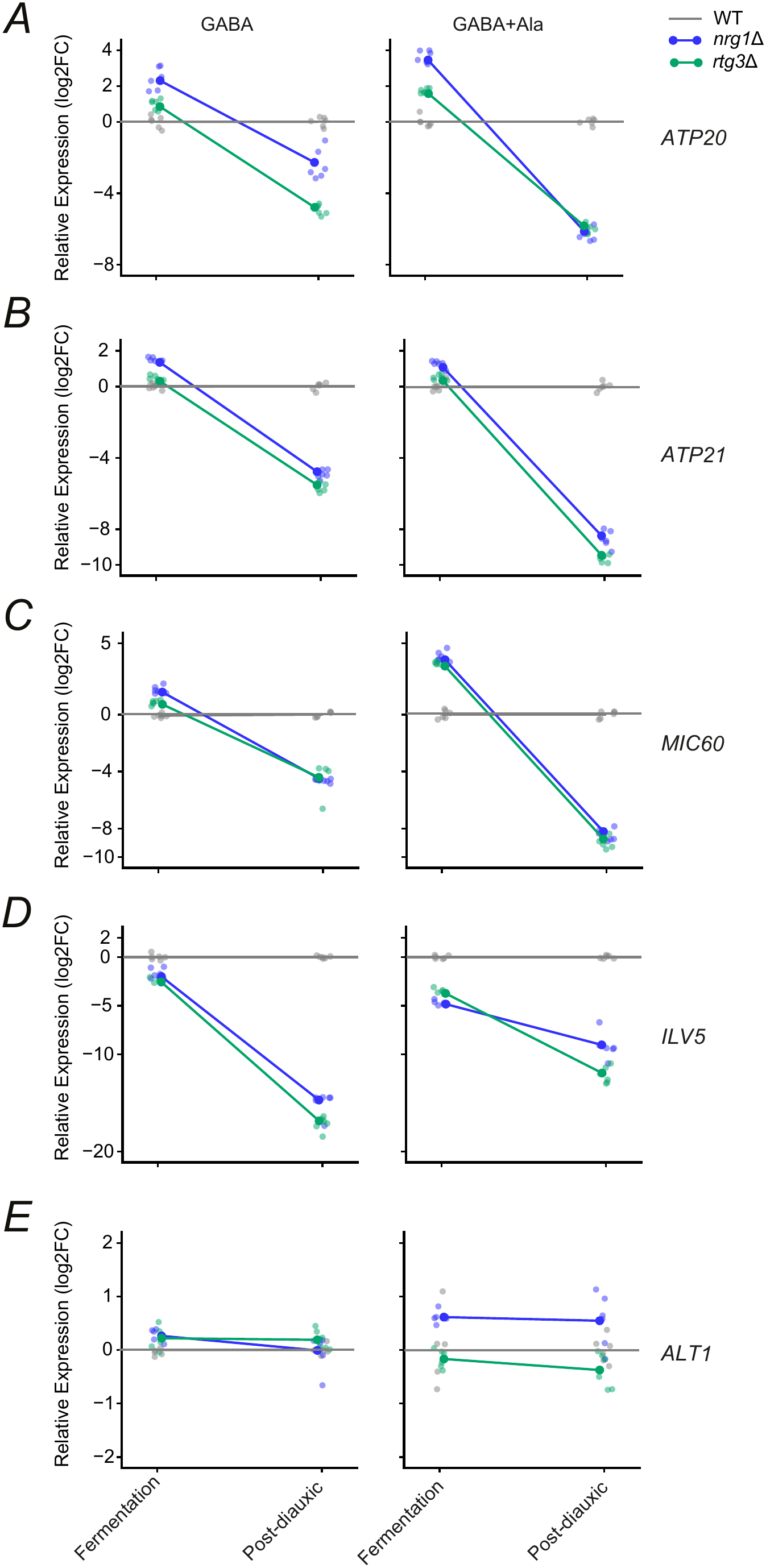
Nrg1-Rtg3 switches from repression to activation of cristae-organizing genes during the diauxic transition. RT-qPCR analysis of **(A)** *ATP20*, **(B)** *ATP21*, **(C)** *MIC60*, selected as key cristae-organizing genes, together with **(D)** *ILV5* as a mitochondrial nucleoid gene regulated by Nrg1-Rtg3 and **(E)** *ALT1* as an alanine-responsive, Nrg1-Rtg3-independent control, in WT, *nrg1*Δ, and *rtg3*Δ cells during fermentative and post-diauxic growth. Cells were grown in minimal medium containing 2% glucose and 7 mM GABA, with or without 7 mM alanine, and collected at OD_600_ = 0.3 (fermentative) or OD_600_ = 1.0 (post-diauxic). Expression was normalized to 18S rRNA and calculated by the 2^−ΔΔCt^ method. For each treatment and growth phase, expression is shown as log2 fold change relative to the corresponding WT sample, which was set to zero. Solid symbols indicate mean values and lighter symbols indicate individual biological replicates; lines connect mean values between growth phases. *ILV5* and *ALT1* were included as controls for Nrg1-Rtg3-dependent and Nrg1-Rtg3-independent expression, respectively.

Consistent with this connection, loss of *ATP20* itself produced pronounced mitochondrial ultrastructural defects whose predominant morphology changed across the diauxic transition. During fermentative growth, *atp20*Δ cells were enriched for irregular mitochondrial profiles and onion-ring structures, whereas after the diauxic transition mitochondria lacking recognizable cristae became the predominant class (**Fig. 6A,B**; **Fig. S5**). Thus, the structural consequences of *ATP20* loss vary with metabolic state, further linking this Nrg1-Rtg3 target to inner-membrane remodeling. Together, these results identify a distinct class of Nrg1-Rtg3 targets whose mode of regulation switches across the diauxic transition. Nrg1-Rtg3 represses cristae-organizing genes during fermentative growth and becomes required for their activation in the post-diauxic state, providing a transcriptional link between metabolic state and mitochondrial cristae remodeling.

**Figure 6.**
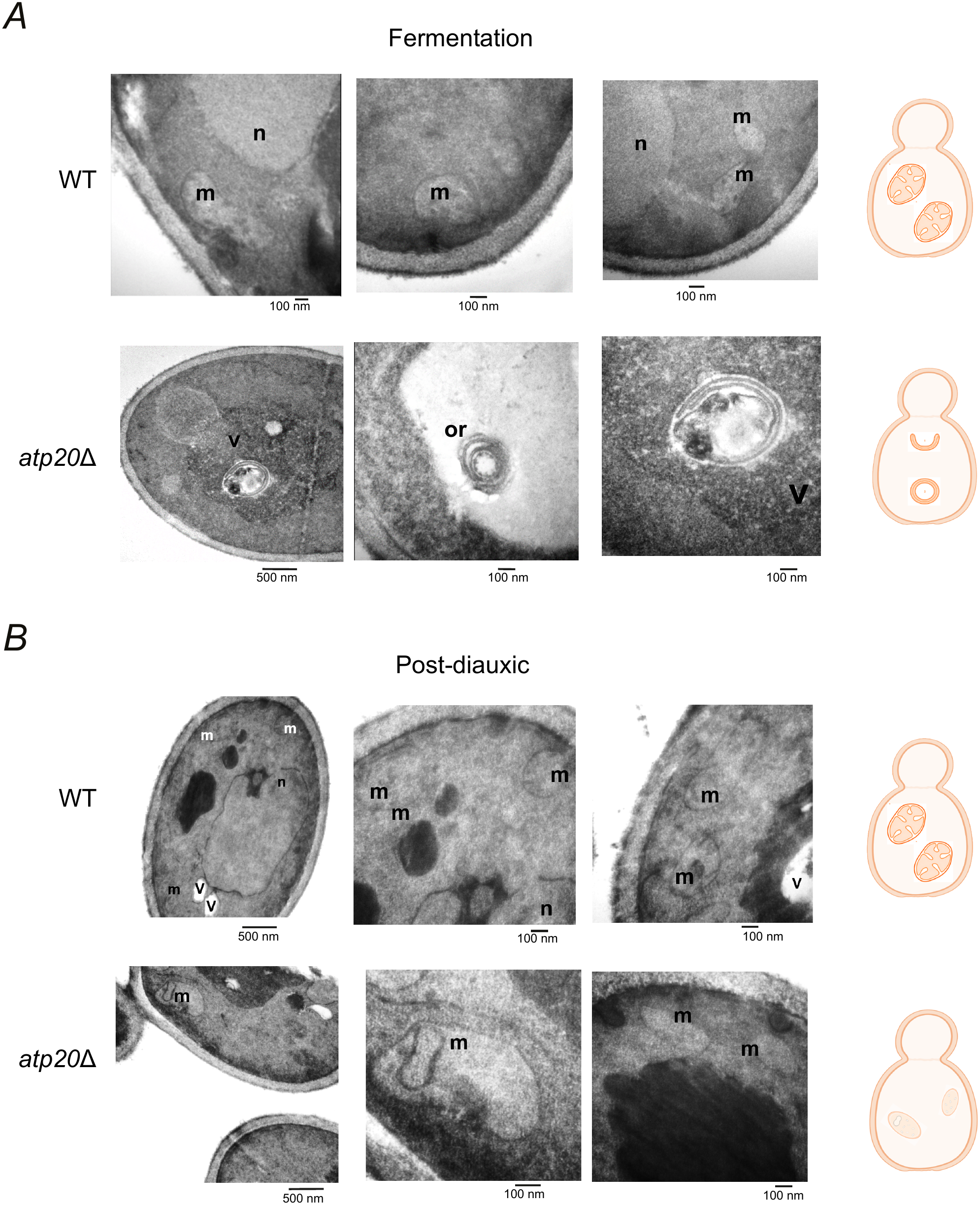
Loss of Atp20 alters mitochondrial cristae architecture during the diauxic transition. Representative TEM images of WT and *atp20*Δ cells (BY4742 background) during **(A)** fermentative growth (OD_600_ = 0.3) and **(B)** post-diauxic growth (OD_600_ = 1.0) in minimal medium containing 2% glucose and 40 mM ammonium sulfate. Three representative images are shown for each strain, with schematics at right summarizing the predominant mitochondrial morphology. During fermentative growth, *atp20*Δ cells frequently displayed onion-ring structures, whereas after the diauxic transition mitochondria lacking recognizable cristae predominated. **n**, nucleus; **m**, mitochondrion; **V**, vacuole; **or**, onion-ring mitochondrial structure. Scale bars are indicated.

### The diauxic transition switches Nrg1-Rtg3 promoter recruitment at *ATP20*

The switch from repression to activation of the cristae-organizing genes raised the question of how Nrg1 and Rtg3 generate opposite transcriptional outputs across the diauxic transition. The nucleoid targets described above provided a reference for comparison. At *ILV5* and *MGM101*, Rtg3 binds independently and is required for Nrg1 recruitment, supporting transcriptional activation (**Fig. 2**). We therefore asked whether the switch in *ATP20* regulation was accompanied by a change in the recruitment relationship between the two factors. ChIP-qPCR across the *ATP20* promoter revealed strong occupancy of both Nrg1 and Rtg3 during fermentative growth, with Nrg1 showing the predominant enrichment across the regulatory region (**Fig. 7A**). Nrg1 occupancy was largely retained in *rtg3*Δ cells, indicating that Nrg1 associates with the promoter independently of Rtg3 during repression (**Fig. 7B**). Rtg3 occupancy, however, was reduced in *nrg1*Δ cells (**Fig. 7C**). Thus, during fermentative growth, recruitment of Rtg3 to *ATP20* depends on Nrg1. This relationship is the reverse of that observed at the activated nucleoid promoters *ILV5* and *MGM101*, where Nrg1 recruitment depends on Rtg3.

**Figure 7.**
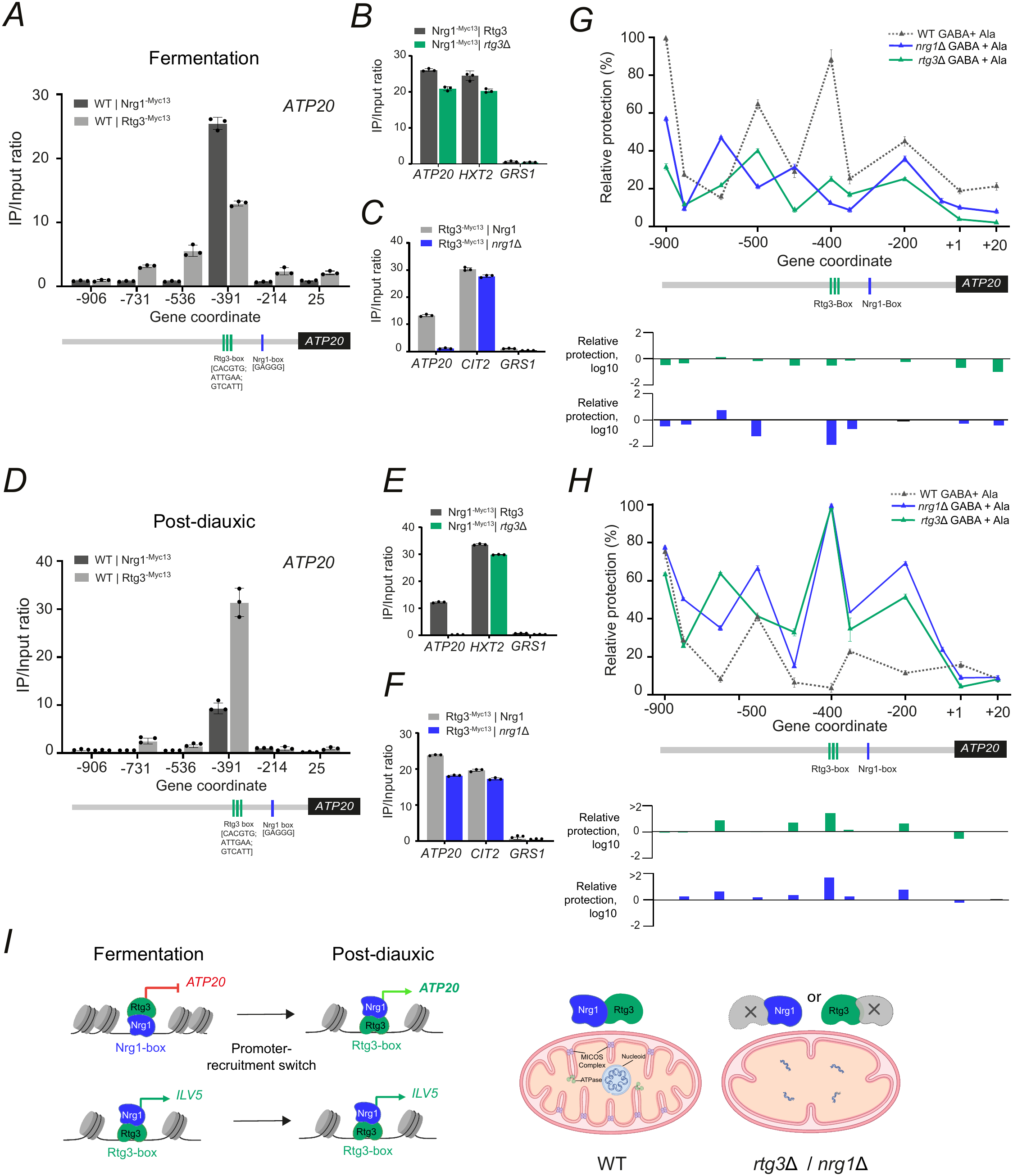
Nrg1-Rtg3 promoter recruitment at *ATP20* reverses across the diauxic transition. (A,D) ChIP-qPCR profiles of Nrg1-Myc13 and Rtg3-Myc13 across the *ATP20* promoter during fermentative growth (**A**; OD_600_ = 0.3) and after the diauxic transition (**D**; OD_600_ = 1.0). Cells were grown in minimal medium containing 2% glucose and 7 mM GABA. Amplicon coordinates are relative to the translation start site; predicted Rtg3-and Nrg1-binding sites are indicated below the promoter schematic. **(B,C,E,F)** ChIP-qPCR analysis of Nrg1-Myc13 in WT and *rtg3*Δ cells **(B,E)** and Rtg3-Myc13 in WT and *nrg1*Δ cells **(C,F)** during fermentative **(B,C)** and post-diauxic **(E,F)** growth. *HXT2* and *CIT2* were included as positive controls for Nrg1 and Rtg3 binding, respectively, and *GRS1* as a negative control. ChIP enrichment is shown as the IP/input ratio. **(G,H)** Nucleosome scanning analysis across the *ATP20* promoter in WT, *nrg1*Δ, and *rtg3*Δ cells grown with GABA plus alanine during fermentative **(G)** and post-diauxic **(H)** growth. Upper plots show relative protection across the promoter; lower plots show the log10 ratio of relative protection in each mutant relative to WT at the corresponding positions. Predicted Rtg3-and Nrg1-binding sites are indicated. Data represent three independent experiments. **(I)** Model summarizing the recruitment switch at *ATP20* and its contrast with *ILV5*. Nrg1-dependent recruitment of Rtg3 at *ATP20* is associated with repression during fermentation, whereas Rtg3-dependent recruitment of Nrg1 is associated with post-diauxic activation. At *ILV5*, Rtg3-dependent recruitment of Nrg1 is maintained across both metabolic states. Mitochondrial schematics summarize the cristae-deficient phenotype associated with loss of either Nrg1 or Rtg3.

We observed that this recruitment relationship was reversed after the diauxic transition. In the post-diauxic state, Rtg3 became the predominant factor detected across the *ATP20* regulatory region, whereas Nrg1 occupancy was reduced relative to the fermentative condition (**Fig. 7D**). More importantly, Nrg1 binding was essentially lost in *rtg3*Δ cells, while Rtg3 occupancy remained substantial in the absence of Nrg1 (**Fig. 7E,F**). Thus, activation of *ATP20* after the diauxic transition coincides with a reversal in recruitment dependence: Rtg3 recruitment depends on Nrg1 during repression, whereas Nrg1 recruitment depends on Rtg3 during activation. We observed the same reversal at *ATP21* and *MIC60*. During fermentative growth, Nrg1 recruitment at both promoters was largely independent of Rtg3, whereas Rtg3 recruitment showed greater dependence on Nrg1. After the diauxic transition, Nrg1 became strongly dependent on Rtg3, while Rtg3 retained promoter occupancy in *nrg1*Δ cells (**Fig. S6**). Thus, reversal of the recruitment relationship accompanies the switch from repression to activation across multiple cristae-organizing genes.

We next asked whether Nrg1 and Rtg3 associate with proteins linked to chromatin regulation. IP-MS was performed separately for each factor, and we focused on proteins recovered in both immunoprecipitates relative to the HA negative control. Gene Ontology enrichment analysis of this shared set highlighted chromatin-related functions: chromatin organization and chromatin remodeling were among the most significant biological-process terms, and chromatin binding was enriched among molecular functions. Enriched cellular-component terms included the SLIK and SAGA complexes, the INO80 complex, and the RSC-type complex, identifying several major chromatin-regulatory complexes among the proteins recovered with both factors (**Fig. S7**; **Dataset S2**). Because these complexes have established roles in transcription-associated nucleosome remodeling, their recovery suggested a possible link between Nrg1-Rtg3 recruitment and promoter chromatin organization. We therefore tested directly whether the recruitment switch at *ATP20* was accompanied by changes in local nucleosome organization.

Nucleosome scanning revealed pronounced differences between the repressed fermentative and activated post-diauxic states (**Fig. 7G,H**). During fermentative growth, the WT promoter displayed several regions of high relative protection across the upstream regulatory sequence, including prominent peaks surrounding the Nrg1-and Rtg3-binding regions. Loss of either factor reduced protection at several of these positions, consistent with the derepression observed in both mutants **(Fig. 7G**). After the diauxic transition, WT cells showed markedly lower protection across much of the central promoter, whereas both *nrg1*Δ and *rtg3*Δ cells showed increased protection over this region (**Fig. 7H**). Thus, although the factor that can bind independently changes across the diauxic transition, both Nrg1 and Rtg3 are required to establish the WT chromatin organization associated with each transcriptional state.

Together, our results support a model in which a switch in Nrg1-Rtg3 promoter recruitment accompanies the transition of *ATP20* from repression to activation (**Fig. 7I**). During fermentative growth, Nrg1 can associate with the *ATP20* promoter independently and is required for efficient Rtg3 recruitment, a configuration associated with repression and the corresponding WT nucleosome organization. After the diauxic transition, this relationship reverses: Rtg3 can bind independently and is required for Nrg1 recruitment as *ATP20* becomes activated and the promoter adopts a different nucleosome organization. In contrast, at the nucleoid-associated target *ILV5*, Rtg3-dependent recruitment of Nrg1 is maintained across both metabolic states and is consistently associated with transcriptional activation. The model therefore illustrates how Nrg1 and Rtg3 can maintain activation of a mitochondrial nucleoid program while changing their recruitment relationship at cristae-organizing genes as cells transition to respiratory growth. Loss of either factor disrupts both regulatory outputs, providing a transcriptional connection to the combined defects in mitochondrial genome maintenance and cristae architecture observed in the mutants.

## Discussion

The metabolic flexibility of *Saccharomyces cerevisiae* requires mitochondria to remain genetically competent during fermentative growth while acquiring the architecture needed for efficient respiration as glucose becomes limiting. Our results place Nrg1-Rtg3 at the intersection of these two requirements, with the heteromeric regulator directly activating the mitochondrial nucleoid genes *ILV5* and *MGM101*, providing a transcriptional link to its previously described role in mitochondrial DNA maintenance [13]. At the same time, loss of either factor produces severe defects in mitochondrial cristae architecture that are already evident during fermentative growth and persist after the diauxic transition. In parallel, Nrg1-Rtg3 represses *ATP20*, *ATP21*, and *MIC60* during fermentation, whereas both factors are required for activation of these genes after the diauxic transition. The corresponding switch in promoter recruitment provides a mechanism through which the same regulator can maintain mitochondrial genome competence while reprogramming cristae organization as metabolic requirements change.

The transition from fermentation to respiration involves extensive changes in transcriptional regulation, including the coordinated action of repressors and activators controlling carbon source utilization and respiratory metabolism [3,4]. Individual regulators can also exert opposing transcriptional effects during this transition. Ert1, for example, contributes to activation of gluconeogenic genes while repressing *PDC1*, a major fermentation gene, during growth on ethanol [4]. Nrg1-Rtg3 constitutes a distinct regulatory mechanism, because its effect on the three cristae-organizing genes switches from repression to activation as cells cross the diauxic transition. This behavior is particularly notable given the established activities of its two components. Nrg1 functions in glucose-dependent repression, including recruitment of the Cyc8-Tup1 corepressor, whereas Rtg3 functions with Rtg1 as a transcriptional activator in the mitochondrial retrograde response [11,12]. Our results show that these activities can be combined within Nrg1-Rtg3 in a metabolically dependent manner, allowing its regulatory output at the same promoters to change from repression during fermentation to activation after the diauxic transition.

The promoter occupancy experiments provide a mechanistic basis for this switch from repression to activation. During fermentative growth, Nrg1 can associate with the *ATP20*, *ATP21*, and *MIC60* promoters independently of Rtg3, whereas Rtg3 recruitment strictly depends on Nrg1, leading to transcriptional repression. After the diauxic transition, this relationship reverses, with Rtg3 displaying independent promoter occupancy and Nrg1 becoming strongly dependent on Rtg3 for recruitment, leading to transcriptional activation. This recruitment pattern resembles the Rtg3-dependent recruitment of Nrg1 observed at the activated *ILV5* and *MGM101* promoters, while contrasting with the Nrg1-dependent recruitment of Rtg3 which leads to *ALT2* repression [13]. These observations suggest that the direction of promoter recruitment dependence between Nrg1 and Rtg3 contributes to the regulatory output of the heteromeric regulator. Nucleosome-scanning experiments further show that both Nrg1 and Rtg3 are required to maintain WT chromatin organization at the *ATP20* promoter, linking the switch in promoter recruitment with changes in promoter chromatin organization. The nature of the signal that drives the described switch from Nrg1-Rtg3 promoter recruitment remains to be determined.

Direct regulation of *ILV5* and *MGM101* also provides a plausible mechanism for the mitochondrial DNA phenotype previously associated with Nrg1-Rtg3 [13]. Mitochondrial DNA is organized into nucleoids whose protein components contribute to genome organization, inheritance, and stability [18]. Mgm101 has a direct role in mitochondrial DNA maintenance and recombination, whereas Ilv5 combines its metabolic function with an additional role in nucleoid organization and mtDNA stability [19–22]. The phenotypes observed here are consistent with these different contributions. Loss of *MGM101* closely reproduced the complete mtDNA depletion and respiratory deficiency of *nrg1*Δ and *rtg3*Δ, whereas *ilv5*Δ cells retained approximately half of the WT mtDNA level and remained capable of slower growth under respiratory conditions. In our experiments, disruption of several other nucleoid components similarly coupled mtDNA loss to respiratory deficiency. *ILV5* and *MGM101* therefore provide direct transcriptional routes through which Nrg1-Rtg3 contributes to mitochondrial genome competence, although the severity of the transcription-factor mutants is unlikely to be explained by the sole activity of these two targets.

Mitochondrial cristae remodeling represents a second, metabolically dynamic output of Nrg1-Rtg3 regulation. During fermentative growth, yeast mitochondria contain relatively sparse cristae, whereas the transition toward respiratory metabolism is accompanied by increased mitochondrial abundance and development of a more extensive inner membrane [14,15]. ATP synthase oligomerization and the MICOS complex are central determinants of this architecture. Atp20 and Atp21 are required for ATP synthase dimerization and normal cristae morphology [16,17], while Mic60 plays a central role in the organization of cristae junctions [17]. Regulation of *ATP20*, *ATP21*, and *MIC60* by Nrg1-Rtg3 therefore provides a direct connection between metabolic state and genes that shape the mitochondrial inner membrane. An important feature of the phenotype is that *nrg1Δ* and *rtg3Δ* cells already show severe cristae defects during fermentative growth, when these genes are repressed in a WT strain. In this context, cristae architecture depends on the coordinated action of ATP synthase oligomers, MICOS, Mgm1, and membrane lipid composition, so altered expression of these three targets contribute to the phenotype but is unlikely to fully explain it [15].

Interpretation of the cristae phenotype must also account for the severe mtDNA depletion in *nrg1*Δ and *rtg3*Δ cells. Both mutants lack detectable mitochondrial DNA and consequently fail to sustain respiratory growth [13]. The relevance of mtDNA depletion to cristae organization is supported by the reported phenotype of cells lacking Mip1, a factor required for mtDNA maintenance. These cells become ρ0 and show severe cristae disorganization, similar to that observed in *nrg1*Δ and *rtg3*Δ cells [23]. Thus, the ultrastructural phenotypes of the Nrg1 and Rtg3 mutants may reflect mtDNA depletion as well as altered nuclear regulation of cristae-organizing genes. At the same time, the transcriptional effects on *ATP20*, *ATP21*, and *MIC60* establish an additional route through which Nrg1-Rtg3 can influence inner-membrane organization. The *atp20*Δ phenotype supports this connection, given that these cells form onion-ring structures and their predominant mitochondrial morphology changes across the diauxic transition. This shows that disruption of a single Nrg1-Rtg3 target can produce pronounced mitochondrial abnormalities. Because the *atp20*Δ phenotype differs from the severe cristae deficiency in *nrg1*Δ and *rtg3*Δ cells, that deficiency may reflect the combined disruption of mtDNA maintenance and transcriptional regulation of several cristae-organizing genes rather than failure of a single structural component.

Our study distinguishes two mitochondrial programs with different requirements across the diauxic transition. Mitochondrial genome competence must be preserved even during fermentative growth, when respiration has a relatively weaker contribution to ATP production, because an intact mitochondrial genome is required when cells subsequently depend on respiratory metabolism [13,15]. Cristae organization, in contrast, changes substantially as respiratory demand increases [14,15]. This distinction is reflected in Nrg1-Rtg3 regulation of its two target classes. At the nucleoid genes *ILV5* and *MGM101*, Rtg3 binds independently and is required for Nrg1 recruitment, while Nrg1-Rtg3-dependent activation of *ILV5* is maintained across metabolic states. By contrast, regulation of the cristae-organizing genes *ATP20*, *ATP21*, and *MIC60* switches from repression during fermentative growth to activation after the diauxic transition, coinciding with a reversal in recruitment dependence from Nrg1-dependent Rtg3 recruitment to Rtg3-dependent Nrg1 recruitment. Together, these findings show that Nrg1-Rtg3 sustains the mitochondrial nucleoid gene program across metabolic states, while recruitment dependence at cristae-organizing genes reverses as their regulation shifts from repression during fermentation to activation after the diauxic transition.

## Methods

The key resources used in this study are listed in **Table S1** and **Dataset S3.**

### Strains and media

*S. cerevisiae* strains used in this study are listed in **Table S1**. The CLA1 strain and the BY4742 strain were used as parental backgrounds. The CLA1 strain and its derivatives were used to investigate the contribution of Nrg1 and Rtg3 in regulation of mitochondrial function and respiratory metabolism. Previously generated nrg1Δ and rtg3Δ derivatives of CLA1 were obtained from [13]. To evaluate the contribution of genes involved in mitochondrial function, *ilv5*Δ, *mgm101*Δ, *rim1*Δ, *abf2*Δ, *aco1*Δ, *atp20*Δ, *atp21*Δ, and *mic6*0Δ strains were analyzed. The corresponding deletion strains in the BY4742 background were obtained from Open Biosystems (YSC1054), whereas the *ilv5*Δ strain in the CLA1 background was generated in this study. Complementation strains were generated by introducing the corresponding wild-type gene on the plasmid pRS416, including pRS416-ILV5 and pRS416-MGM101 (**Table S1**). To examine the relation between Nrg1 and Rtg3, strains carrying epitope-tagged versions of these transcriptional regulators were used, including Nrg1^-Myc13^ and Rtg3^-TAP^ [13]

Strains were grown on minimal medium (MM) containing salts, trace elements, and vitamins according to the formula for yeast nitrogen base (Difco). Glucose (2% w/v) or ethanol (2% v/v) was used as the carbon source; 40 mM ammonium sulfate was used as nitrogen source. Uracil (20 mg/L), leucine (100 mg/L), histidine (20 mg/L), lysine (30 mg/L), valine (150 mg/L), isoleucine (30 mg/L) or glutamate (20 mg/L) were added as auxotrophic requirements when needed. Cells were incubated at 30 °C with shaking (250 rpm).

### Construction of *S. cerevisiae ilv5*Δ strain

The single *ilv5*Δ mutant strain was constructed as follows: The *ILV5* (SGD:S000004347) gene was replaced by homologous recombination in strain *S. cerevisiae* CLA1 using a module containing the NatMX cassette flanked by 60 bp of 5‘upstream region and 60 bp of 3‘UTR sequence of *ILV5.* This module was amplified from p4339 plasmid [24] by PCR with A1 and A2 deoxyoligonucleotides. The PCR product was transformed into the *S. cerevisiae* CLA1 strain. Transformants were selected for nourseothricin resistance (100 µg/mL).

### Construction of pRS416-*ILV5* and pRS416-*MGM101* plasmids

*ILV5* and *MGM101* gene sequence was amplified, including their 5‘and 3‘UTR sequences and cloned into plasmid pRS416 (*CEN6 ARSH4 URA3*) [25]. The PCR products were digested, ligated and cloned to generate pRS416-*ILV5* and pRS416-*MGM101*, which were transformed into strain CLA 1-1 *(ilv5*Δ) and BY4742-1 *(mgm101*Δ). Transformants were selected for uracil prototrophy on minimal medium. The *ILV5* gene was amplified with deoxyoligonucleotides Clon1 and Clon2. The insert included the 5‘upstream region comprised of 760 bp (from-760 to the start codon), and a 3‘UTR from the stop codon to +461 bp. The *MGM101* gene was amplified with deoxyoligonucleotides Clon3 and Clon4. In this case, the 5‘UTR region comprised from-826 bp to the start codon and the 3‘UTR from the stop codon to +389. Both PCR products and pRS416 plasmid were digested with *XhoI-BamHI* restriction enzymes and ligated after purification. Ligations were transformed into *DH5*α *Escherichia coli* strain. The plasmids pRS416-*ILV5* and pRS416-*MGM101* were purified from *E. coli* extracts after cloning and transformed into *S. cerevisiae* strains.

### RNA-seq and differential expression analysis

*S. cerevisiae* strains were grown on MM glucose (2% w/v) as a carbon source and supplemented with GABA (7 mM) or GABA + Ala (7 mM) as nitrogen sources to an OD_600_ of 0.3–0.5. The cells were harvested by centrifugation at 3000 rpm. for 5 min and washed twice with distilled H_2_O + DEPC. Four independent biological replicates were made per condition. The cells were centrifuged and the pellet was resuspended in 1.0 mL of AE buffer (50 mM sodium acetate and 10 mM EDTA, pH 5.2). Total RNA was extracted using a modified phenol-SDS method. Briefly, 40 µL of 10% SDS, 450 µL of acid phenol, and approximately 400 µL of glass beads (0.45 mm diameter) were added to the cell suspension. The mixture was vigorously vortexed and incubated at 65°C for 20 min, with intermittent vortexing for 30 s every 5 min. It was then rapidly frozen in liquid nitrogen for 3 min. Samples were centrifuged at 10,000 rpm (6,700 × g) for 10 min at 4°C in a centrifuge to separate phases, and the aqueous phase was recovered and purified with two sequential extractions using phenol/chloroform/isoamyl alcohol (25:24:1), followed by a third extraction with chloroform/isoamyl alcohol (24:1). RNA was precipitated with 0.3 M sodium acetate and 2.5 volumes of absolute ethanol, washed with 70% ethanol, air-dried at room temperature, and resuspended in nuclease-free water.

Total RNA was purified from collected samples as described above and precipitated with 7.5 M LiCl following [26]. RNA quality and integrity were assessed by 1% denaturing agarose gel electrophoresis, spectrophotometry (NanoDrop, Thermo Scientific), and analysis using a Bioanalyzer 2100 (Agilent Technologies). Only samples with OD260/280 and OD260/230 ratios above 2.0 and RNA integrity number (RIN) values ≥ 4.0 were used for sequencing.

RNA samples were sent to Novogene Co. (Sacramento, CA, USA) for RNA library construction and sequencing. Poly(A)-enriched mRNA was isolated using oligo(dT) magnetic beads, fragmented, and reverse-transcribed using random hexamer primers. Library construction included end repair, adapter ligation, size selection (∼150 bp), PCR amplification, and quality control using Qubit and RT-PCR. Paired-end sequencing (≥ 20 million read pairs per sample) was performed on the Illumina HiSeq 6000 platform. Bioinformatic processing was carried out using the snakePipes (v3.0.0) mRNA-Seq pipeline [27] with established default parameters. In brief, the quality of raw reads and alignments was evaluated using FastQC and MultiQC [28]. Raw paired-end reads were processed with Cutadapt [29] to remove Illumina adapters, low-quality bases (Phred score < 16), and ambiguous bases (N) from the 3′ end of each read, with a minimum read length of 25 bases. Filtered reads were then aligned to the *S. cerevisiae* R64-1-1 genome from Ensembl (corresponding to NCBI GCA_000146045.2) using STAR [30], and gene-level read counts were generated with featureCounts [31]. Differential gene expression analysis was performed using DESeq2 [32]. Transcript abundance was expressed as transcripts per million (TPM) and used for experimental quality control.

Functional enrichment analysis of differentially expressed genes was conducted using FungiFun webtool [33] and annotations from the Kyoto Encyclopedia of Genes and Genomes (KEGG) database. Enrichment was considered significant at an adjusted *p* value < 0.05. KEGG pathway visualization was performed using the KEGG REST API [34].

### RT-qPCR analysis of gene expression

*Saccharomyces cerevisiae* strains were grown to the indicated OD_600_ on MM glucose (2% w/v) as carbon source and supplemented with 40 mM ammonium sulfate, GABA (7 mM) or GABA + Ala (7 mM) as nitrogen sources to an OD_600_ of 0.3-0.5 or 1.0-1.2. Cells were harvested by centrifugation at 3000 rpm for 5 min and washed twice with distilled H_2_O. The cells were then transferred to a mortar, frozen with liquid nitrogen and mechanically disrupted with the aid of a pestle. Total RNA was extracted using TRIzol reagent (Invitrogen) following the TDS procedure. The RNA quality and concentration were determined using a NanoDrop 2000 spectrophotometer (NanoDrop Technologies). RNA was treated with RQ1 RNAse-Free DNAse (Promega) to degrade DNA from the samples; afterwards, cDNA was obtained using the RevertAid H Minus First Strand cDNA Synthesis Kit, following the manufacturer instructions (Thermo Scientific). cDNA samples were adjusted to a concentration of 30 ng µl^-1^. PCR was performed using the KAPA SYBR Fast kit (Roche) and run on the Corbett Research Rotor-Gene 6000 (Qiagen). Gene expression was obtained for *ALT1, ILV5, MGM101, RIM1, ACO1, ABF2, COX6, COX2, COX3, ATP6, ATP9, COB1, MIC60, ATP20 and ATP21,* using deoxyoligonucleotides QC1 to QC32. Data were analyzed using the 2^—ΔΔCT^ method, in which the RDN18S (18S) constitutive gene was used as reference gene [35]. Statistical analysis was performed using GraphPad Prism 9.0 (GraphPad Software Inc). All experiments were repeated three times, and the results are expressed as mean ± SEM.

### Quantification of mtDNA content by qPCR

To quantify the mtDNA, we obtained the amount of mtDNA relative to nDNA. We followed the method described in [13], estimating the mtDNA/nDNA ratio by qPCR using different mtDNA genes (*COX2*, *COX3*, and *ATP6*) and, as a nuclear-encoded gene, *COX6*, using deoxyoligonucleotides QC3 to QC8 and QC13 to QC14. Total DNA from the indicated strains was extracted twice by phenol/chloroform/isoamyl alcohol (25:24:1). Samples were incubated with 20 μg RNase A for 15 min at 37°C. DNA precipitation was carried out with an equal volume of ethanol for 30 min at −20°C and resuspended in nuclease-free water. DNA was quantified by NanoDrop (Thermo Scientific), and samples were diluted to 50 ng μl^−1^ for qPCR assay. qPCR was performed using SYBR Green dye (2X KAPA SYBR FAST; Roche). Conditions for qPCR were 94°C for 5 min (one cycle), 94°C for 15 s, 59°C for 30 s, and 72°C for 20 s (30 cycles). We obtained the cycle threshold (Ct) for each sample, and following the 2^—ΔΔCT^ method, we calculated the mtDNA/nDNA ratio as described previously [36].

### Nucleosome scanning assay

The nucleosome scanning assay (NuSA) was performed as previously described [37]. Cultures (100 mL) containing glucose (2% w/v) as a carbon source and supplemented with GABA (7 mM) or GABA (7 mM) + Ala (7 mM) as nitrogen sources were grown at OD_600nm_ = 0.3–0.5 or 1.0–1.2. To determine nucleosome positioning at the *ILV5, MGM101, RIM1,* and *ATP20* promoters, samples were processed as previously described [38,39]. Quantitative PCR (qPCR) analysis was performed using a Corbett Life Science Rotor Gene 6000 and SYBR Green dye (2X KAPA SYBR FAST, Invitrogen). Real-time PCR was performed as follows: 94°C for 5 min (1 cycle), 94°C for 15 s, 58°C for 20 s, and 72°C for 20 s (35 cycles). The PCR deoxyoligonucleotides (ILV5_1 to ILV5_27) used for *ILV5,* (MGM101_1 to MGM101_17) for *MGM101*, (RIM1_1 to RIM1_18) for *RIM1* and (ATP20_1 to ATP20_6) for *ATP20* analysis are described in supplementary material. Relative protection was calculated as a ratio to the control of *VCX1*, which was amplified using QC33 and QC34 deoxyoligonucleotides.

### Quantitative chromatin immunoprecipitation (qChIP)

Formaldehyde cross-linking and immunoprecipitation were performed according to a previously described procedure [40]. Yeast cells (100 mL of OD_600nm_ = 0.3–0.5 or 1.0–1.2) were cross-linked with 1% formaldehyde for 20 min at room temperature. Afterwards, 125 mM glycine was added, and the mixture was incubated for 5 min. The cells were harvested and washed with PBS. The pelleted cells were suspended in lysis buffer (140 mM NaCl, 1 mM EDTA, 50 mM HEPES/KOH, 1% Triton X-100, and 0.1% sodium deoxycholate) with a protease inhibitor cocktail (Complete Mini, Roche). The cells were lysed with glass beads and collected by centrifugation. Extracts were sonicated using a Diagenode Bioruptor to produce chromatin fragments with an average size of 300 bp.

Immunoprecipitation reactions were carried out with 1 µg anti-c-Myc antibody (9E11, Santa Cruz Biotechnology) and protein A beads for 3 h, washed, suspended in TE buffer/1% SDS, and incubated overnight at 65°C to reverse the formaldehyde cross-linking. Immunoprecipitates were then incubated with proteinase K (Roche), followed by phenol/chloroform/isoamyl alcohol extraction, precipitation, and suspension in 30 μl of TE buffer. Dilutions of input DNA (1:100) and immunoprecipitated DNA (1:2) were analyzed by qPCR. Real-time PCR-based DNA amplification was performed using primers that were initially screened for dimer absence or cross-hybridization. Only the deoxyoligonucleotides (ChILV_1 to ChILV_12) used for *ILV5,* (ChMGM_1 to ChMGM_12) for *MGM101*, (ChRIM1_1 to ChRIM1_12) for *RIM1*, (ChATP20_1 to ChATP20_12) used for *ATP20*, (ChATP21_1 to ChATP21_10) used for *ATP21* and (ChMIC60_1 to ChMIC60_14) used for *MIC60* with similar amplification efficiencies were used.

Quantitative chromatin immunoprecipitation (qChIP) analysis was performed using a Corbett Life Science Rotor-Gene 6000 instrument. For each qPCR-amplified region, the ratio of immunoprecipitated DNA to total input DNA was calculated as IP/input = 2^(Input Ct − IP Ct) [41]. The results represent the mean values and SEM of at least three independent cross-linked samples, each immunoprecipitated twice with the antibody. The *HXT2* gene regulated by Nrg1 amplified with ChHXT2_1 and ChHXT2_2 deoxyoligonucleotides, and the *CIT2* gene regulated by Rtg3 amplified with ChCIT2_1 and ChCIT2_2 deoxyoligonucleotides were used as positive controls for DNA binding in the qChIP assay. The *GRS1* gene amplified with ChGRS_1 and ChGRS_2 deoxyoligonucleotides was used as a negative control.

### Immunoprecipitation of Rtg3-TAP and Nrg1-Myc13 for proteomic analysis

Immunoprecipitation was performed using a modified version of the method described by [42]. *S. cerevisiae* strains were grown in MM containing glucose (2% w/v) as the carbon source and supplemented with GABA (7 mM) or GABA (7 mM) plus alanine (7 mM). Cells were harvested at OD_600_ = 0.5 by centrifugation at 3,000 rpm for 5 min and washed with 1× TBS. Cell pellets were transferred to 2-mL microcentrifuge tubes and frozen in liquid nitrogen for 15 s. Pellets were resuspended in 500 μl lysis buffer (50 mM HEPES, 200 mM NaOAc, 0.1 mM EDTA, 0.1 mM EGTA, 5 mM MgOAc, 5% glycerol, 0.25% NP-40, 3 mM DTT, 1 mM PMSF, and protease inhibitor cocktail), and 400 μl of 0.5-mm glass beads were added. Cells were disrupted by vortexing twice for 4 min, with a 2-min incubation on ice between cycles. Lysates were centrifuged at 14,000 rpm for 5 min at 4°C, and the supernatant containing the total protein extract was transferred to a new 2-mL microcentrifuge tube.

Protein concentration was determined using the Bradford assay. For each immunoprecipitation, 2 mg of total protein was incubated with 60 μl Pierce protein A/G agarose beads (Thermo Scientific) and 10 μl of TAP antibody CAB1001 (Invitrogen) or anti-Myc antibody 9E11 (Santa Cruz Biotechnology). Samples were incubated with gentle agitation for 2 h at 4°C. Beads were then washed five times with 1 mL wash buffer (50 mM HEPES, 200 mM NaOAc, 0.1 mM EDTA, 0.1 mM EGTA, 5 mM MgOAc, 5% glycerol, 0.25% NP-40, 3 mM DTT, and 1 mM PMSF), with centrifugation at 2,000 rpm for 1 min at 4°C between washes. After the final wash, the supernatant was completely removed and 50 μl loading buffer (50 mM Tris, pH 6.8, 2% SDS, 10% glycerol, 100 mM DTT, and 1 mM PMSF) was added. Samples were incubated at 65°C for 10 min and centrifuged at 2,000 rpm for 3 min at 4°C. Forty microliters of the resulting supernatant were used for proteomic analysis.

### Sample preparation and tryptic digestion for LC-MS/MS

Samples were adjusted to pH 8.6 with 3 M Tris base (pH 12) and reduced with dithiothreitol (DTT) at 56°C for 1 h. Samples were then alkylated with 50 mM iodoacetamide (IAM, Sigma-Aldrich) for 1 h at room temperature in the dark. Proteins were subsequently precipitated by adding nine volumes of absolute ethanol and incubating overnight at −20°C. Precipitated proteins were pelleted by centrifugation at 16,000 x g for 15 min at 4°C. The pellet was washed twice with 90% ethanol. Proteins were digested with 1 µg of mass spectrometry-grade trypsin (Promega) per sample in 50 mM ammonium bicarbonate (Sigma-Aldrich) for 16–18 h at 37°C. After digestion, peptides were desalted, dried to completion, and stored at −80°C until LC-MS/MS analysis.

### LC-MS/MS data acquisition

For LC-MS/MS analysis, peptides were solubilized in 0.1% formic acid (mobile phase A) and analyzed using a Vanquish Neo nano-liquid chromatography system (Thermo Fisher Scientific) coupled to a Q Exactive Plus mass spectrometer (Thermo Fisher Scientific). Peptides were first desalted using a PepMap Neo C18 trap cartridge (5 μm, 300 μm x 5 mm; Thermo Fisher Scientific) and subsequently separated on an EASY-Spray PepMap Neo C18 analytical column (2 μm, 75 μm × 500 mm, 1500 bar; Thermo Fisher Scientific) at a flow rate of 300 nL/min.

Chromatographic separation was performed using a gradient of mobile phase B (0.1% formic acid in 80% acetonitrile): 6–20% B for 60 min, 20–35% B for 30 min, 35–99% B for 1 min, and 99% B for 9 min. For data-independent acquisition (DIA), 32 isolation windows spanning 390–1200 m/z with a 1-Da overlap were used. Resolution was set to 70,000 for MS1 and 35,000 for MS2, with a normalized collision energy (NCE) of 30%.

### Protein identification and statistical analysis

Raw files were processed using DIA-NN version 1.8.1 [43]. Search parameters included methionine oxidation and N-terminal acetylation as variable modifications, with up to one missed cleavage and one variable modification allowed per peptide. Match-between-runs (MBR) was enabled, and Robust LC (High Precision) was used for quantification. The false discovery rate (FDR) was set to 1%. Proteins were retained when at least two peptides were identified and an intensity value was obtained in at least two of three biological replicates.

Protein abundance was compared using the one-factor Statistical Analysis module of MetaboAnalyst 6.0 [44]. Pairwise comparisons were performed using volcano plots with median normalization, log10 transformation, no filtering, and no data scaling. Thresholds were set at a 1.5-fold change and *p* < 0.05. Putative Nrg1-Rtg3 interactors were defined as proteins enriched in both Nrg1-Myc13 and Rtg3-TAP immunoprecipitates relative to the HA control according to these criteria. Functional enrichment analysis of the putative interactors was performed using DAVID Bioinformatics Resources [45].

### Transmission electron microscopy

Transmission electron microscopy was performed as previously described [46,47], with minor modifications. Yeast cells were grown in minimal medium (MM) containing 2% glucose and 40 mM ammonium sulfate. Cells were harvested at the indicated growth phases and fixed overnight at 4°C in 4% glutaraldehyde prepared in 0.2 M PIPES buffer (pH 6.8) supplemented with 0.2 M sorbitol, 2 mM MgCl₂, and 2 mM CaCl₂. After fixation, samples were washed with distilled water and post-fixed with 2% potassium permanganate for 45 min at room temperature. Cells were then stained with 1% uranyl acetate for 1 h and dehydrated through a graded ethanol series from 30% to 100%.

Following dehydration, samples were infiltrated with Durcupan™ ACM resin and embedded for polymerization. Ultrathin sections (60 nm) were obtained using a Leica Ultracut UCT ultramicrotome and mounted on Formvar-coated copper grids. Sections were subsequently stained with 0.35% uranyl acetate and 4% lead citrate and examined using a JEOL 1010 transmission electron microscope operated at 80 kV. Digital micrographs were acquired using a Hamamatsu camera (Hamamatsu Photonics K.K., Japan).

## Data availability

Supplementary datasets supporting this study are provided with the manuscript. RNA-seq data have been deposited in the Gene Expression Omnibus (GEO) repository under accession number GSE351438. Proteomic data have been deposited in the iProX integrated proteome resources repository under accession number IPX0019651001.

## Author contributions

Conceptualization: CS, AG. Methodology: AV-A, CC-B, ER, VG-M, NT-R, RO-H, HQ, JG. Formal analysis: FP-G, ACP-H, EM-M, ALT-M, SE-G, NT-R, RO-H, XM-E, HQ, JG, LR-R, LP-R, AD, CS. Investigation: AV-A, CC-B, ER, FP-G, ACP-H, VG-M, ALT-M, SE-G, NT-R, RO-H, XM-E, HQ, JG, LR-R, LP-R. Resources: ACP-H, SE-G, NT-R, AG. Data curation: FP-G, ACP-H, SE-G, HQ, LP-R. Writing – original draft: AG. Writing – review & editing: AV-A, CC-B, AD, CS. Visualization: AV-A, CC-B, EM-M, JG, AD. Supervision: XM-E, LR-R, AD, CS, AG. Project administration: CC-B, AG. Funding acquisition: AG.

## Acknowledgements

We are grateful to Manuel L. Robert for fruitful and motivating discussions, which deeply influenced our work, and to Hugo Hernández for engaging discussions. We are grateful to Laura Ongay, Guadalupe Códiz, and Minerva Mora from the Molecular Biology Unit at IFC, UNAM, to Carlos Alberto Peralta Alvarez from the Bioinformatics Unit at IFC, UNAM, and to Magdalena Hernández-Ortiz from Centro de Ciencias Genómicas, UNAM, for work with proteomic analysis. We are also indebted to Rocio Romualdo Martínez for helpful secretarial assistance. During the preparation of this manuscript, the authors used ChatGPT GPT-6 (OpenAI) to assist with English-language editing and improve clarity. The authors reviewed and edited the output as needed and take full responsibility for the content of the published article.

## Funding

This study was performed in partial fulfillment of the requirements for A.V-A and V.G.-M their master’s degrees in Biochemical Sciences at the Universidad Nacional Autónoma de México, which were carried with a fellowship from the Secretaría de Ciencia, Humanidades, Tecnología e Innovación (SECIHTI, CVU 2113097 and CVU 2053664, respectively). This study was funded by the Dirección General de Asuntos del Personal Académico (DGAPA), Universidad Nacional Autónoma de México (UNAM), grants IN207424 and IN203926 (http://dgapa.unam.mx), and grant CBF-2025-1-962 from SECIHTI. Edgar Ramírez held a SECIHTI Posdoctoral Program fellowship (CVU 472824).

## Competing interest

The authors declare no competing interests.

## Supplemental Material

**Figure S1:** Summary of RNA-seq differential expression and pathway enrichment analyses.

**Figure S2:** Differential regulation of mitochondrial nucleoid genes by Nrg1 and Rtg3.

**Figure S3:** Rtg3 has the predominant role in chromatin organization at the *RIM1* promoter.

**Figure S4:** Mitochondrial gene expression, respiratory growth, and mtDNA maintenance in Nrg1-Rtg3 and nucleoid mutants.

**Figure S5:** Quantification of mitochondrial phenotypes during fermentative and post-diauxic growth.

**Figure S6:** Nrg1-Rtg3 promoter recruitment at *ATP21* and *MIC60* reverses across the diauxic transition.

**Figure S7:** Functional enrichment of proteins associated with both Nrg1 and Rtg3.

**Table S1:** Biological materials, reagents, and tools used in this study.

**Dataset S1:** RNA-seq expression data and differential-expression results for wild-type, *nrg1Δ*, and *rtg3Δ* cells.

**Dataset S2:** Mass spectrometry data for proteins recovered in Nrg1^-Myc13^, Rtg3^-TAP^, and HA-control immunoprecipitations.

**Dataset S3:** Complete list of deoxyoligonucleotides used in this study.

## Notes

### Competing Interest Statement

The authors have declared no competing interest.

### Summary of Updates

This revised version 2 includes all Supplementary Material

## References

1. Dashko S, Zhou N, Compagno C, Piškur J. Why, when, and how did yeast evolve alcoholic fermentation? FEMS Yeast Research. 2014;14: 826–832. doi:10.1111/1567-1364.12161

2. DeRisi JL, Iyer VR, Brown PO. Exploring the metabolic and genetic control of gene expression on a genomic scale. Science. 1997;278: 680–686. doi:10.1126/science.278.5338.680

3. Turcotte B, Liang XB, Robert F, Soontorngun N. Transcriptional regulation of nonfermentable carbon utilization in budding yeast. FEMS Yeast Research. 2010;10: 2–13. doi:10.1111/j.1567-1364.2009.00555.x

4. Gasmi N, Jacques P-E, Klimova N, Guo X, Ricciardi A, Robert F, et al. The Switch from Fermentation to Respiration in Saccharomyces cerevisiae Is Regulated by the Ert1 Transcriptional Activator/Repressor. Genetics. 2014;198: 547–560. doi:10.1534/genetics.114.168609

5. Hedbacker K, Carlson M. SNF1/AMPK pathways in yeast. Frontiers in Bioscience. 2008;13: 2408–2420. doi:10.2741/2854

6. Buschlen S, Amillet J, Guiard B, Fournier A, Marcireau C, Bolotin-Fukuhara M. The S. cerevisiae HAP complex, a key regulator of mitochondrial function, coordinates nuclear and mitochondrial gene expression. Comparative and Functional Genomics. 2003;4: 37–46. doi:10.1002/cfg.254

7. Young ET, Dombek KM, Tachibana C, Ideker T. Multiple pathways are co-regulated by the protein kinase Snf1 and the transcription factors Adr1 and Cat8. Journal of Biological Chemistry. 2003;278: 26146–26158. doi:10.1074/jbc.M301981200

8. Soontorngun N, Larochelle M, Drouin S, Robert F, Turcotte B. Regulation of Gluconeogenesis in Saccharomyces cerevisiae Is Mediated by Activator and Repressor Functions of Rds2. Molecular and Cellular Biology. 2007;27: 7895–7905. doi:10.1128/MCB.01055-07

9. Proft M, Struhl K. Hog1 kinase converts the Sko1-Cyc8-Tup1 repressor complex into an activator that recruits SAGA and SWI/SNF in response to osmotic stress. Molecular Cell. 2002;9: 1307–1317. doi:10.1016/s1097-2765(02)00557-9

10. Gsaller F, Hortschansky P, Beattie SR, Klammer V, Tuppatsch K, Lechner BE, et al. The Janus transcription factor HapX controls fungal adaptation to both iron starvation and iron excess. The EMBO Journal. 2014;33: 2261–2276. doi:10.15252/embj.201489468

11. Park SH, Koh SS, Chun JH, Hwang HJ, Kang HS. Nrg1 Is a Transcriptional Repressor for Glucose Repression of STA1 Gene Expression in Saccharomyces cerevisiae. Molecular and Cellular Biology. 1999;19: 2044–2050. doi:10.1128/MCB.19.3.2044

12. Liu Z, Butow RA. Mitochondrial retrograde signaling. Annual Review of Genetics. 2006;40: 159–185. doi:10.1146/annurev.genet.40.110405.090613

13. Campero-Basaldua C, González J, García JA, Ramírez E, Hernández H, Aguirre B, et al. Neo-functionalization in Saccharomyces cerevisiae: a novel Nrg1-Rtg3 chimeric transcriptional modulator is essential to maintain mitochondrial DNA integrity. Royal Society Open Science. 2023;10. doi:10.1098/rsos.231209

14. Di Bartolomeo F, Malina C, Campbell K, Mormino M, Fuchs J, Vorontsov E, et al. Absolute yeast mitochondrial proteome quantification reveals trade-off between biosynthesis and energy generation during diauxic shift. Proceedings of the National Academy of Sciences. 2020;117: 7524–7535. doi:10.1073/pnas.1918216117

15. Klecker T, Westermann B. Pathways shaping the mitochondrial inner membrane. Open Biology. 2021;11: 210238. doi:10.1098/rsob.210238

16. Paumard P, Vaillier J, Coulary B, Schaeffer J, Soubannier V, Mueller DM, et al. The ATP synthase is involved in generating mitochondrial cristae morphology. The EMBO Journal. 2002;21: 221–230. doi:10.1093/emboj/21.3.221

17. Rabl R, Soubannier V, Scholz R, Vogel F, Mendl N, Vasiljev-Neumeyer A, et al. Formation of cristae and crista junctions in mitochondria depends on antagonism between Fcj1 and Su e/g. The Journal of Cell Biology. 2009;185: 1047–1063. doi:10.1083/jcb.200811099

18. Miyakawa I. Organization and dynamics of yeast mitochondrial nucleoids. Proceedings of the Japan Academy, Series B. 2017;93: 339–359. doi:10.2183/pjab.93.021

19. Zelenaya-Troitskaya O, Perlman PS, Butow RA. An enzyme in yeast mitochondria that catalyzes a step in branched-chain amino acid biosynthesis also functions in mitochondrial DNA stability. The EMBO Journal. 1995;14: 3268–3276. doi:10.1002/j.1460-2075.1995.tb07330.x

20. Bateman JM, Iacovino M, Perlman PS, Butow RA. Mitochondrial DNA Instability Mutants of the Bifunctional Protein Ilv5p Have Altered Organization in Mitochondria and Are Targeted for Degradation by Hsp78 and the Pim1p Protease. Journal of Biological Chemistry. 2002;277: 47946–47953. doi:10.1074/jbc.M209071200

21. Meeusen S, Tieu Q, Wong E, Weiss E, Schieltz D, Yates JR, et al. Mgm101p Is a Novel Component of the Mitochondrial Nucleoid That Binds DNA and Is Required for the Repair of Oxidatively Damaged Mitochondrial DNA. The Journal of Cell Biology. 1999;145: 291–304. doi:10.1083/jcb.145.2.291

22. Mbantenkhu M, Wang X, Nardozzi JD, Wilkens S, Hoffman E, Patel A, et al. Mgm101 Is a Rad52-related Protein Required for Mitochondrial DNA Recombination. Journal of Biological Chemistry. 2011;286: 42360–42370. doi:10.1074/jbc.M111.307512

23. Guerrero-Castillo S, van Strien J, Brandt U, Arnold S. Ablation of mitochondrial DNA results in widespread remodeling of the mitochondrial complexome. The EMBO Journal. 2021;40: e108648. doi:10.15252/embj.2021108648

24. Goldstein AL, McCusker JH. Three new dominant drug resistance cassettes for gene disruption in Saccharomyces cerevisiae. Yeast. 1999;15: 1541–1553. doi:10.1002/(SICI)1097-0061(199910)15:14<1541::AID-YEA476>3.0.CO;2-K

25. Sikorski RS, Hieter P. A system of shuttle vectors and yeast host strains designed for efficient manipulation of DNA in Saccharomyces cerevisiae. Genetics. 1989;122: 19–27. doi:10.1093/genetics/122.1.19

26. Walker SE, Lorsch J. RNA Purification – Precipitation Methods. Methods in Enzymology. 2013;530: 337–343. doi:10.1016/B978-0-12-420037-1.00019-1

27. Bhardwaj V, Heyne S, Sikora K, Rabbani L, Rauer M, Kilpert F, et al. snakePipes: facilitating flexible, scalable and integrative epigenomic analysis. Bioinformatics. 2019;35: 4757–4759. doi:10.1093/bioinformatics/btz436

28. Ewels P, Magnusson M, Lundin S, Käller M. MultiQC: summarize analysis results for multiple tools and samples in a single report. Bioinformatics. 2016;32: 3047–3048. doi:10.1093/bioinformatics/btw354

29. Martin M. Cutadapt removes adapter sequences from high-throughput sequencing reads. EMBnet.journal. 2011;17: 10–10. doi:10.14806/ej.17.1.200

30. Dobin A, Davis CA, Schlesinger F, Drenkow J, Zaleski C, Jha S, et al. STAR: ultrafast universal RNA-seq aligner. Bioinformatics. 2013;29: 15–21. doi:10.1093/bioinformatics/bts635

31. Liao Y, Smyth GK, Shi W. featureCounts: an efficient general purpose program for assigning sequence reads to genomic features. Bioinformatics. 2014;30: 923–930. doi:10.1093/bioinformatics/btt656

32. Love MI, Huber W, Anders S. Moderated estimation of fold change and dispersion for RNA-seq data with DESeq2. Genome Biology. 2014;15: 550. doi:10.1186/s13059-014-0550-8

33. Priebe S, Linde J, Albrecht D, Guthke R, Brakhage AA. FungiFun: A web-based application for functional categorization of fungal genes and proteins. Fungal Genetics and Biology. 2011;48: 353–358. doi:10.1016/j.fgb.2010.11.001

34. Kanehisa M, Furumichi M, Sato Y, Matsuura Y, Ishiguro-Watanabe M. KEGG: biological systems database as a model of the real world. Nucleic Acids Research. 2025;53: D672–D677. doi:10.1093/nar/gkae909

35. Jozefczuk J, Adjaye J. Quantitative Real-Time PCR-Based Analysis of Gene Expression. Methods in Enzymology. 2011;500: 99–109. doi:10.1016/B978-0-12-385118-5.00006-2

36. Quiros PM, Goyal A, Jha P, Auwerx J. Analysis of mtDNA/nDNA Ratio in Mice. Current Protocols in Mouse Biology. 2017;7: 47–54. doi:10.1002/cpmo.21

37. Infante JJ, Law GL, Young ET. Analysis of Nucleosome Positioning Using a Nucleosome-Scanning Assay. Methods in Molecular Biology. 2012;833: 63–87. doi:10.1007/978-1-61779-477-3_5

38. Campero-Basaldua C, Quezada H, Riego-Ruíz L, Márquez D, Rojas E, González J, et al. Diversification of the kinetic properties of yeast NADP-glutamate-dehydrogenase isozymes proceeds independently of their evolutionary origin. MicrobiologyOpen. 2017;6. doi:10.1002/mbo3.419

39. González J, López G, Argueta S, Escalera-Fanjul X, el Hafidi M, Campero-Basaldua C, et al. Diversification of Transcriptional Regulation Determines Subfunctionalization of Paralogous Branched Chain Aminotransferases in the Yeast Saccharomyces cerevisiae. Genetics. 2017;207: 975–991. doi:10.1534/genetics.117.300290

40. Hernández H, Aranda C, López G, Riego L, González A. Hap2-3-5-Gln3 determine transcriptional activation of GDH1 and ASN1 under repressive nitrogen conditions in the yeast Saccharomyces cerevisiae. Microbiology. 2011;157: 879–889. doi:10.1099/mic.0.044974-0

41. Litt MD, Simpson M, Gaszner M, Allis CD, Felsenfeld G. Correlation Between Histone Lysine Methylation and Developmental Changes at the Chicken β-Globin Locus. Science. 2001;293: 2453–2455. doi:10.1126/science.1064413

42. Gerace E, Moazed D. Coimmunoprecipitation of Proteins from Yeast. Methods in Enzymology. 2014;541: 13–26. doi:10.1016/B978-0-12-420119-4.00002-1

43. Demichev V, Messner CB, Vernardis SI, Lilley KS, Ralser M. DIA-NN: neural networks and interference correction enable deep proteome coverage in high throughput. Nature Methods. 2020;17: 41–44. doi:10.1038/s41592-019-0638-x

44. Pang Z, Lu Y, Zhou G, Hui F, Xu L, Viau C, et al. MetaboAnalyst 6.0: towards a unified platform for metabolomics data processing, analysis and interpretation. Nucleic Acids Research. 2024;52: W398–W406. doi:10.1093/nar/gkae253

45. Sherman BT, Hao M, Qiu J, Jiao X, Baseler MW, Lane HC, et al. DAVID: a web server for functional enrichment analysis and functional annotation of gene lists (2021 update). Nucleic Acids Research. 2022;50: W216–W221. doi:10.1093/nar/gkac194

46. Unger A-K, Geimer S, Harner M, Neupert W, Westermann B. Analysis of Yeast Mitochondria by Electron Microscopy. Methods in Molecular Biology. 2017;1567: 293–314. doi:10.1007/978-1-4939-6824-4_18

47. Rosas-Paz M, Zamora-Bello A, Torres-Ramírez N, Villarreal-Huerta D, Romero-Aguilar L, Pardo JP, et al. Nitrogen limitation-induced adaptive response and lipogenesis in the Antarctic yeast Rhodotorula mucilaginosa M94C9. Frontiers in Microbiology. 2024;15: 1416155. doi:10.3389/fmicb.2024.1416155

