## Supplementary material for "A switch in Nrg1-Rtg3 promoter recruitment coordinates mitochondrial cristae remodeling during the diauxic transition": Table S1

**Table S1. Biological materials, reagents, and tools used in this study.**

| REAGENT or RESOURCE | SOURCE | IDENTIFIER |
|---|---|---|
| Antibodies | | |
| Ac α TAP | Invitrogen | CAB1001 |
| Ac α MYC (9E11) | Santa Cruz Biotechnology | Sc-47694 |
| Ac α MYC (9E10) | Santa Cruz Biotechnology | Sc-40 |
| Ac α HA | Santa Cruz Biotechnology | Sc-7392 |
| Ac α anti-rabbit | Santa Cruz Biotechnology | Sc-2357 |
| Bacterial strains | | |
| *E. coli* DH5 *α* | This paper | N/A |
| Chemicals, peptides, and recombinant proteins | | |
| Protease Inhibitor Cocktail | Complete Mini, Roche | 11697498001 |
| Tri reagent solution | Invitrogen | AM9738 |
| RNAse-Free DNAse | Promega | M6101 |
| Proteinase K Solution | Roche | 25530-049 |
| Geneticin (G418) | Gibco | 10463982 |
| Neursotricine (Nat) | Jena Bioscience | 96736-11-7 |
| XhoI restriction enzyme | NEB | R0146S |
| BamHI restriction enzyme | NEB | R0136S |
| Dynbeads Protein A | Invitrogen / Thermo Fisher Scientific | 10006D |
| Pierce protein A/G agarose beads | Thermo Scientific | 20421 |
| Iodoacetamide | Merck Sigma-Aldrich | I1149-25G |
| Mass spectrometry grade trypsin | Promega | V5280 |
| Durcupan™ ACM | Sigma-Aldrich | 44611 |
| Critical commercial assays | | |
| PCRBIO HiFi Polymerase | PCR BIOSYSTEMS | PB10.41-02 |
| Rapid DNA ligation Kit | Roche | 11635379001 |
| Shrimp Alkaline Phosphatase | New England Biolabs | M0371S |
| QIAprep Spin Miniprep kit | Qiagen | 27106 |
| Wizard SV gel and PCR clean-up system | Promega | A9282 |
| RevertAid H Minus First Strand cDNA Synthesis Kit | Thermo Scientific | K1622 |
| qPCR GreenMaster highROX | Jena Bioscience | PCR-374L |
| Deposited data | | |
| RNAseq data | This paper | GEO GSE351438 (sncnckmydtipncn) |
| Proteomics data | This paper | iProX IPX0019651001 |
| *S. cerevisiae* R64-1-1 reference genome | Ensembl | NCBI GCA_000146045.2 |
| *S. cerevisiae ILV5* gene sequence | Saccharomyces Genome Database | SGD: S000004347 |
| Experimental models: strains | | |

| | | |
|---|---|---|
| CLA 1 : MAT$\alpha$ *ILV5 MGM101 ACO1 ABF2 RIM1 ATP20 ATP21 MIC60 NRG1 RTG3* ura3$^{-}$ leu2$^{-}$ | Campero-Basaldua et al. 2023 | N/A |
| CLA 1-2 (*nrg1*Δ): MAT$\alpha$ *nrg1*Δ::KanMX4 *ILV5 MGM101 ACO1 ABF2 RIM1 ATP20 ATP21 MIC60 RTG3* ura3$^{-}$ leu2$^{-}$ | Campero-Basaldua et al. 2023 | N/A |
| CLA 1-2 (*rtg3* Δ): MAT$\alpha$ *rtg3*Δ::NatMX4 *ILV5 MGM101 ACO1 ABF2 RIM1 ATP20 ATP21 MIC60 NRG1* ura3$^{-}$ leu2$^{-}$ | Campero-Basaldua et al. 2023 | N/A |
| CLA1 (pRS416) MAT$\alpha$/pRS416 *ILV5 MGM101 ACO1 ABF2 RIM1 ATP20 ATP21 MIC60 NRG1 RTG3* leu2$^{-}$ | This study | N/A |
| CLA 1-1 *(ilv5*Δ): MATα *ilv5*Δ::NatMX4 *MGM101 ACO1 ABF2 RIM1 ATP20 ATP21 MIC60 NRG1 RTG3*ura3$^{-}$ leu2$^{-}$ | This study | N/A |
| CLA 1-1 *(ilv5*Δ / pRS416) MAT$\alpha$/ pRS416 *ilv5*Δ::NatMX4 *MGM101 ACO1 ABF2 RIM1 ATP20 ATP21 MIC60 NRG1 RTG3* leu2$^{-}$ | This study | N/A |
| CLA 1-1 *(ilv5*Δ / pRS416- *ILV5):* MAT$\alpha$ *ilv5*Δ::NatMX4 / pRS416- *ILV5 MGM101 ACO1 ABF2 RIM1 ATP20 ATP21 MIC60 NRG1 RTG3* leu2$^{-}$ | This study | N/A |
| BY4742: MAT$\alpha$ *ILV5 MGM101 ACO1 ABF2 RIM1 ATP20 ATP21 MIC60 NRG1 RTG3* ura3$^{-}$ leu2$^{-}$ lys2$^{-}$ His3$^{-}$ | Open Biosystems | YSC1054 |
| BY4742 (pRS416): MAT$\alpha$ *ILV5 MGM101 ACO1 ABF2 RIM1 ATP20 ATP21 MIC60 NRG1 RTG3* leu2$^{-}$ lys2$^{-}$ His3$^{-}$ | This study | N/A |
| BY4742-1 *(mgm101*Δ*):* MAT$\alpha$ *mgm101*Δ::KanMX4 *ILV5 ILV5 ACO1 ABF2 RIM1 ATP20 ATP21 MIC60 NRG1 RTG3* ura3$^{-}$ leu2$^{-}$ lys2$^{-}$ His3$^{-}$ | Open Biosystems | YSC1054 |
| BY4742-1 *(mgm101*Δ */* pRS416*):* MAT$\alpha$ *mgm101*Δ::KanMX4 *ILV5 ACO1 ABF2 RIM1 ATP20 ATP21 MIC60 NRG1 RTG3* leu2$^{-}$ lys2$^{-}$ His3$^{-}$ | This study | N/A |
| BY4742-1 *(mgm101*Δ */* pRS416-*MGM101):* MAT$\alpha$ *mgm101*Δ::KanMX4 / pRS416-*MGM101 ILV5 ACO1 ABF2 RIM1 ATP20 ATP21 MIC60 NRG1 RTG3* leu2$^{-}$ lys2$^{-}$ His3$^{-}$ | This study | N/A |
| BY4742-2 (*rim1*Δ*):* MAT$\alpha$ *rim1*Δ::KanMX4 *ILV5 MGM101 ACO1 ABF2 ATP20 ATP21 MIC60 NRG1 RTG3* ura3$^{-}$ leu2$^{-}$ lys2$^{-}$ His3$^{-}$ | Open Biosystems | YSC1054 |
| BY4742-3 (*abf2*Δ*):* MAT$\alpha$ *abf2*Δ::KanMX4 *ILV5 MGM101 ACO1 RIM1 ATP20 ATP21 MIC60 NRG1 RTG3* ura3$^{-}$ leu2$^{-}$ lys2$^{-}$ His3$^{-}$ | Open Biosystems | YSC1054 |
| BY4742-4 (*aco1*Δ*):* MAT$\alpha$ *aco1*Δ::KanMX4 *ILV5 MGM101 ABF2 RIM1 ATP20 ATP21 MIC60 NRG1 RTG3* ura3$^{-}$ leu2$^{-}$ lys2$^{-}$ His3$^{-}$ | Open Biosystems | YSC1054 |

| | | |
|---|---|---|
| BY4742 (*atp20Δ*) MAT$\alpha$ *atp20*Δ::KanMX4 *ILV5 MGM101 ACO1 ABF2 RIM1 ATP21 MIC60 NRG1 RTG3* ura3$^{-}$ leu2$^{-}$ lys2$^{-}$ his3$^{-}$ | Open Biosystems | YSC1054 |
| BY4742 (*atp21Δ*) MAT$\alpha$ *atp21*Δ::KanMX4 *ILV5 MGM101 ACO1 ABF2 RIM1 ATP20 MIC60 NRG1 RTG3* ura3$^{-}$ leu2$^{-}$ lys2$^{-}$ his3$^{-}$ | Open Biosystems | YSC1054 |
| BY4742 (*mic60Δ*) MAT$\alpha$ *mic60*Δ::KanMX4 *ILV5 MGM101 ACO1 ABF2 RIM1 ATP20 ATP21 NRG1 RTG3* ura3$^{-}$ leu2$^{-}$ lys2$^{-}$ his3$^{-}$ | Open Biosystems | YSC1054 |
| CLA 1-2 (Nrg1$^{-Myc13}$): MAT$\alpha$ Nrg1::$^{MYC13\text{-}KanMX4}$ *ILV5 MGM101 ACO1 ABF2 RIM1 ATP20 ATP21 MIC60 RTG3* ura3$^{-}$ leu2$^{-}$ | Campero-Basaldua et al. 2023 | N/A |
| CLA 1-2 (Nrg1$^{-Myc13}$, *rtg3*Δ): MAT$\alpha$ Nrg1$^{-Myc13::KanMx4}$ *rtg3*Δ::NatMX4 *ILV5 MGM101 ACO1 ABF2 RIM1 ATP20 ATP21 MIC60* ura3 leu2 | Campero-Basaldua et al. 2023 | N/A |
| BY4742 (Rtg3$^{-Tap}$): MAT$\alpha$ Rtg3::$^{-Tap::HIS3}$*ILV5 MGM101 ACO1 ABF2 RIM1 ATP20 ATP21 MIC60 NRG1* ura3$^{-}$ leu2$^{-}$ lys2$^{-}$ | Campero-Basaldua et al. 2023 | N/A |
| BY4742 (Rtg3$^{-Tap}$, *nrg1*Δ): MAT$\alpha$ Rtg3::$^{-Tap::HIS3}$ *nrg1*Δ::KanMX4 *ILV5 MGM101 ACO1 ABF2 RIM1 ATP20 ATP21 MIC60* ura3$^{-}$ leu2$^{-}$ lys2$^{-}$ | Campero-Basaldua et al. 2023 | N/A |
| BY4742 (Rtg3$^{-Tap}$, Nrg1$^{-Myc13}$): MAT$\alpha$ Rtg3::$^{-Tap::HIS3}$ Nrg1$^{-Myc13::KanMx4}$ *ILV5 MGM101 ACO1 ABF2 RIM1 ATP20 ATP21 MIC60* ura3$^{-}$ leu2$^{-}$ lys2$^{-}$ | Campero-Basaldua et al. 2023 | N/A |
| Oligonucleotides | | |
| Primers used to construct null mutants (Table S2, XLSX file) | This paper | N/A |
| Primers used to *ILV5* or *MGM101* cloning (Table S2a, XLSX file) | This paper | N/A |
| Primers used to qPCR assays (Table S2, XLSX file) | This paper | N/A |
| Primers used for ChIP of *ILV5* assay (Table S2, XLSX file) | This paper | N/A |
| Primers used for ChIP of *MGM101* assay (Table S2, XLSX file) | This paper | N/A |
| Primers used for ChIP of *RIM1* assay (Table S2, XLSX file) | This paper | N/A |
| Primers used for ChIP of ATP20 assay (Table S2, XLSX file) | This paper | N/A |
| Primers used for ChIP of *ATP21* assay (Table S2, XLSX file) | This paper | N/A |
| Primers used for ChIP of *MIC60* assay (Table S2, XLSX file) | This paper | N/A |
| Primers used for *ILV5* Nucleosome Scanning Assay ( NuSA) (Table S2, XLSX file) | This paper | N/A |
| Primers used for *MGM101* Nucleosome Scanning Assay ( NuSA) (Table S2, XLSX file) | This paper | N/A |

| | | |
|---|---|---|
| Primers used for *RIM1* Nucleosome Scanning Assay ( NuSA) (Table S2a, XLSX file) | This paper | N/A |
| Primers used for *ATP20* Nucleosome Scanning Assay ( NuSA) (Table S2, XLSX file) | This paper | N/A |
| Recombinant DNA | | |
| Plasmid: pFA6a-kanMX4 | Longtine et al., 1998 | Addgene plasmid 39296 |
| Plasmid: pFA6a-NatMX4 | Goldstein et al., 1999 | Addgene plasmid 134428 |
| Plasmid: pRS416 | Sikorski and Hieter, 1989 | ATCC 87521 |
| Plasmid: pRS416-*ILV5* | This paper | N/A |
| Plasmid: pRS416-*MGM101* | This paper | N/A |
| Software and algorithms | | |
| SnakePipes version 3.0.0 | SnakePipes | https://snakepipes.readthedocs.io/en/stable/ |
| FastQC version 0.12.0 | FastQC | https://github.com/s-andrews/FastQC |
| MultiQC version 1.19 | MultiQC | https://github.com/MultiQC/MultiQC |
| Cutadapt version 4.6 | Martin, 2011 | https://cutadapt.readthedocs.io/en/v4.6/ |
| STAR version 2.7.10b | Dobin et al., 2013 | https://alexdobin.github.io/STAR/ |
| featureCounts (Subread) version 2.0.1 | Liao et al., 2013 | https://subread.sourceforge.net/featureCounts.html |
| R version 4.3.3 | R Core Team | https://www.r-project.org/ |
| DESeq2 version 1.42.0 | Love et al., 2014 | https://bioconductor.org/packages/release/bioc/html/DESeq2.html |
| FungiFun webtool | Priebe et al., 2016 | https://fungifun3.hki-jena.de/ |
| Kyoto Encyclopedia of Genes and Genomes (KEGG) database | KEGG Consortium | https://www.kegg.jp/ |
| CLUSTALW2 | Larkin et al., 2007 | http://www.ebi.ac.uk/Tools/clustalw2 |
| Fungal Genome Search | Fungal Genome Search | https://www.yeastgenome.org/blast-fungal |
| YEASTRACT+ | Teixeira et al., 2023 | https://www.yeastract.com/formseqretrieval.php |
| GraphPad Prism 9.0 | GraphPad Software Inc | https://www.graphpad.com/features |
| DIA-NN program version 1.8.1 | DIA-NN | https://aptila.bio/ |

| | | |
|---|---|---|
| MetaboAnalyst 6.0 program | Pang et al., 2024 | https://www.metaboanalyst.ca/ |
| DAVID Bioinformatics Resources web site | Sherman et al., 2022 | https://davidbioinformatics.nih.gov/home.jsp |
| Other resources | | |
| Nanodrop 2000 | Thermo Scientific | N/A |
| Bioanalyzer 2100 | Agilent Technologies | N/A |
| HiSeq 600 platform | Novogene | Co. (Sacramento, CA, USA) |
| Corbett Research Rotor-Gene 6000 | Corbett Life Science | N/A |
| Sonics vibra cell | Sonics and Materials Inc | N/A |
| Vanquish Neo nano liquid chromatography system | Thermo Fisher Scientific | N/A |
| Q-Exactive Plus mass spectrometer | Thermo Fisher Scientific | N/A |
| PepMap Neo 5 µm C18 300 µm x 5 mm trap cartridge | Thermo Fisher Scientific | 174500 |
| PepMap Neo C18, 2 µm, 75 µm x 500 mm, 1500 bar | Thermo Fisher Scientific | ES75500PN |
| Leica Ultracut UCT ultramicrotome | Leica Biosystems | N/A |
| JEM 1010 transmission electron microscope | Jeol | N/A |
| Hamamatsu camera | Hamamatsu Photonics | N/A |
