## Supplementary Figures S1-S7 for "A switch in Nrg1-Rtg3 promoter recruitment coordinates mitochondrial cristae remodeling during the diauxic transition"

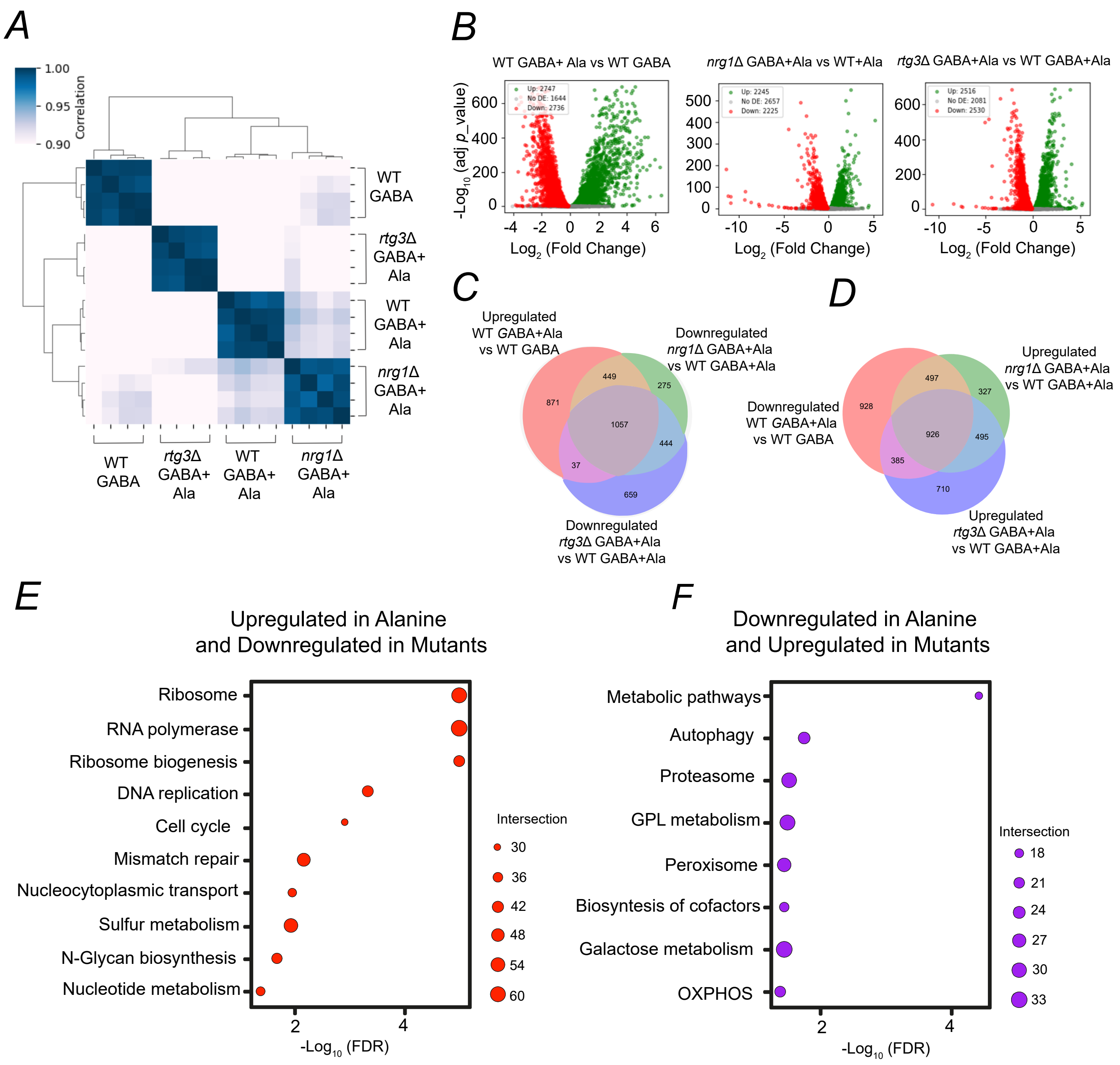


**Figure S1. Summary of RNA-seq differential expression and pathway enrichment analyses. (A)** Sample-to-sample gene expression Pearson correlation heatmap for the 16 RNA-seq samples, with hierarchical clustering. Samples cluster according to their respective experimental groups. **(B)** Volcano plots showing differential gene expression for the following contrasts: WT grown in GABA + alanine versus WT grown in GABA; *nrg1Δ* grown in GABA + alanine versus WT grown in GABA + alanine; and *rtg3Δ* grown in GABA + alanine versus WT grown in GABA + alanine. Each point represents one gene; the x-axis shows log2 fold change and the y-axis shows −log10 BH-adjusted *P* value. Green and red indicate significantly up- and downregulated genes, respectively (adjusted $p < 0.05$). **(C)** Venn diagram showing the overlap among genes upregulated by alanine in WT cells and genes downregulated in *nrg1Δ* and *rtg3Δ* cells relative to WT cells grown in GABA + alanine. **(D)** Venn diagram showing the reciprocal comparison: genes downregulated by alanine in WT cells and genes upregulated in *nrg1Δ* and *rtg3Δ* cells relative to WT cells grown in GABA + alanine. **(E)** Top 10 KEGG pathways enriched among genes in the three-way intersection shown in C. The x-axis indicates −log10(FDR), and point size indicates the number of genes from the intersecting set represented in each pathway. **(F)** KEGG pathways significantly enriched among genes in the three-way intersection shown in D (FDR < 0.05). Axis definitions and point-size encoding are as in E.
